# Precise tuning of cortical contractility regulates cell shape during cytokinesis

**DOI:** 10.1101/635615

**Authors:** Nilay Taneja, Matthew R. Bersi, Sophie Baillargeon, Aidan M. Fenix, James A. Cooper, Ryoma Ohi, Vivian Gama, W. David Merryman, Dylan T. Burnette

## Abstract

The mechanical properties of the cellular cortex regulate shape changes during cell division, cell migration and tissue morphogenesis. During cell division, contractile force generated by the molecular motor myosin II (MII) at the equatorial cortex drives cleavage furrow ingression. Cleavage furrow ingression in turn increases stresses at the polar cortex, where contractility must be regulated to maintain cell shape during cytokinesis. How polar cortex contractility controls cell shape is poorly understood. We show a balance between MII paralogs allows a fine-tuning of cortex tension at the polar cortex to maintain cell shape during cytokinesis, with MIIA driving cleavage furrow ingression and bleb formation, and MIIB serving as a stabilizing motor and mediating completion of cytokinesis. As the majority of non-muscle contractile systems are cortical, this tuning mechanism will likely be applicable to numerous processes driven by MII contractility.

## INTRODUCTION

The cortex of animal cells is composed of a thin network of actin filaments underneath the plasma membrane. The function of the cortex is to provide mechanical support to the membrane and to organize protein complexes (Fritzsche et al., 2016; Salbreux et al., 2012; Sezgin et al., 2017). Membrane and actin cross-linkers, in addition to the molecular motor myosin II (MII), organize and remodel this dynamic structure. Through the modulation of its mechanical properties, the cortex allows cells to change shape and generate tension in response to internal and external physical stimuli. In the case of migratory responses, such as those observed during development and disease progression, the cortex can remodel to produce protrusions such as filopodia, lamellipodia and blebs (Charras and Paluch, 2008; Gardel et al., 2010). During mitosis, the cortex undergoes a series of remodeling events, where initially the cortex allows a cell to round up and then finally divide into two, through the constriction of an equatorial contractile ring (Stewart et al., 2011; Surcel et al., 2010). The cytokinetic phase of cell division presents a major challenge to cortical stability due to the extensive remodeling events required at the cleavage furrow.

During cytokinesis, a transient enrichment of MII and actin at the equatorial cortex creates a contractile network that contains highly organized actomyosin ensembles (Fenix et al., 2016; Fishkind and Wang, 1993; Fujiwara and Pollard, 1976). The polar cortex, however, retains the same isotropic organization found in rounded cells and contains lower MII activity (Bovellan et al., 2014; Levayer and Lecuit, 2012). Accumulation of MII at the cortex creates cortical tension, that in turn, increases intracellular pressure (Stewart et al., 2011). While most studies heavily focus on mechanisms driving contractile ring formation and ingression, relatively fewer studies have studied the polar actomyosin network.

Perturbation of the polar cortical network has been shown to result in shape instability and cytokinetic failure (Guha et al., 2005; Sedzinski et al., 2011). For instance, knockdown of anillin, an actin binding scaffold protein results in a global increase in cortical instability due to enhanced cortical contractility (Sedzinski et al., 2011; Straight et al., 2005). Reducing polar contractility through local delivery of blebbistatin results in cytokinetic failure and misplaced cleavage furrows (Guha et al., 2005). This has resulted in the general idea that the cortex of dividing cells is inherently unstable and prone to shape instabilities, and cortical contractility therefore must be tightly regulated. In this study, we define the role of two MII paralogs, MIIA and MIIB, in driving fine-tuning of polar cortex tension that allows cells to maintain shape during cytokinesis. These findings also uncover a general mechanism that may allow cells to attain a broad range of contractile states in order to perform various cellular functions requiring cortex contractility.

## RESULTS

### MIIA templates MII filament ensembles at the equatorial cortex to drive ingression

We first investigated the role of the predominant MII paralogs in the two cortical networks during cytokinesis. We previously reported that HeLa cells express MIIA and MIIB, but not MIIC (Taneja and Burnette, 2019). MIIA and MIIB filaments are known to co-assemble in the same filaments in the cleavage furrow (Beach et al., 2014). Furthermore, MIIA is required to template MIIB filament stacks at the leading edge of migrating cells during interphase (Fenix et al., 2016). To test whether MIIA is also required for MIIB stack formation in the cleavage furrow, we depleted MIIA using siRNA (MIIA^lo^ cells, Supplementary Fig. 1A-D). We observed a nearly 50% compensatory increase in total MIIB protein levels in MIIA^lo^ cells (Supplementary Fig. 1A).

We used a method we previously developed to measure the length of MII filament stacks using structured illumination microscopy (Fenix et al., 2016). Despite the increase in overall MIIB protein levels, depletion of MIIA significantly reduced the length of MIIB stacks at the cleavage furrow (Fig. 1A, C). Protein level compensation by MIIA upon MIIB depletion (MIIB^lo^ cells) was more modest (∼20%, Supplementary Fig. 1A). Consistent with the templating hypothesis, we found no change in MIIA stacks in the cleavage furrow (Fig. 1B, C). Since there were smaller MII stacks in the cleavage furrow of MIIA^lo^ cells, we hypothesized they would have defects in furrow ingression.

**Figure 1.**
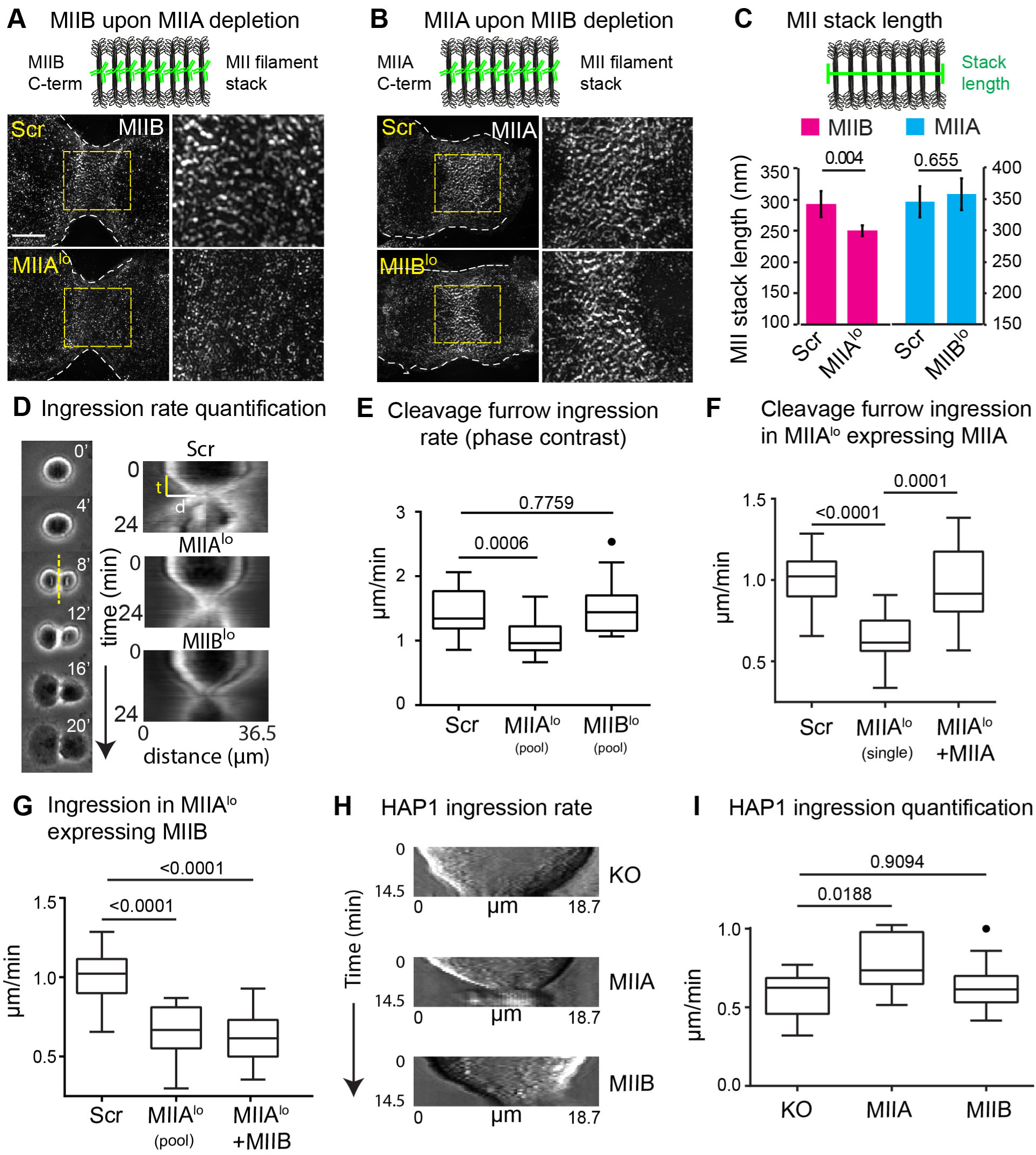
MIIA drives furrow ingression during cytokinesis. A-B) Endogenous MIIB (A) and MIIA (B) in the cleavage furrow of Scr versus MIIA^lo^ or MIIB^lo^ cells, respectively. Insets: enlarged view of yellow box. Schematic shows labeling strategy. C) Average length of MII filament stacks. For (A), n=13 Scr cells over 4 independent experiments; 17 MIIA^lo^ cells over 5 independent experiments. For (B), n=14 Scr and 8 MIIB^lo^ over 3 independent experiments each. Schematic shows labeling strategy. D) Cleavage furrow ingression rate measurement from phase contrast time montages. E) Tukey plots of ingression measured using phase contrast. n=18 Scr, 22 MIIA^lo^ (pooled siRNA) and 20 MIIB^lo^ (pooled siRNA) cells over 3 independent experiments. F) Tukey plots of ingression measured independently using DIC. n=18 Scr, 16 MIIA^lo^ (single siRNA) and 14 MIIA^lo^+MIIA cells over 4 independent experiments. G) Tukey plots of ingression measured independently using DIC. n=15 MIIA^lo^ (pooled siRNA) and 15 MIIA^lo^+MIIB cells over 3 independent experiments. The Scr dataset is the same as (F) and is only displayed for comparison. (H) Representative kymographs of control KO, and MIIA or MIIB expressing HAP1 MIIA KO fibroblasts. (I) Tukey plots of ingression measured using DIC. n= 11 control KO, 12 MIIA expressing and 10 MIIB expressing cells over 3 independent experiments. Solid circles represent outliers. Scale bars- 5 µm.

Measurement of cleavage furrow ingression rates using kymographs generated from phase contrast movies revealed that MIIA depletion indeed caused a significant reduction in cleavage furrow ingression rates compared to scrambled control siRNA (Scr) or MIIB^lo^ cells (Fig. 1D, E). To rule out off-target effects of siRNA knockdown of MIIA, we rescued MIIA depletion using full length siRNA resistant MIIA-mEGFP. Expression of MIIA-mEGFP at 65% of endogenous MIIA levels increased ingression to control rates (Fig. 1F, Table S1). Importantly, overexpression of MIIB at 143% of endogenous MIIB (Table S1) did not increase rate of cleavage furrow ingression in MIIA^lo^ cells (Fig. 1G). This suggested that MIIA is required to drive cleavage furrow ingression. To test whether MIIA is sufficient to increase cleavage furrow ingression rates, we used HAP1 MIIA knockout fibroblasts (Taneja and Burnette, 2019). Expression of MIIA mEGFP, but not MIIB mEGFP increased cleavage furrow ingression rates in HAP1 MIIA KO fibroblasts (Fig. 1H, I). Interestingly, we also found that the amount of MIIA localized to the equatorial cortex positively correlated with ingression rates in human embryonic stem cell (hESC) colonies, that show a natural gradient of MIIA, but not MIIB levels within the colony (Supplementary Fig. 2).

The cortical defects associated with MIIA loss slightly reduced cytokinetic fidelity, resulting in a 3-fold increase in binucleated cells (Fig. 2A-B). As an intended control, we also measured binucleation in MIIB^lo^ HeLa cells. Surprisingly, we observed a dramatic, nearly 10-fold increase in binucleated MIIB^lo^ cells, relative to Scr cells, despite no change in furrow ingression (Fig. 2A-B, Fig. 1E). The increase in binucleation was specific to MIIB depletion, since expression of siRNA resistant MIIB resulted in a significant decrease in binucleation (Fig. 2C).

**Figure 2.**
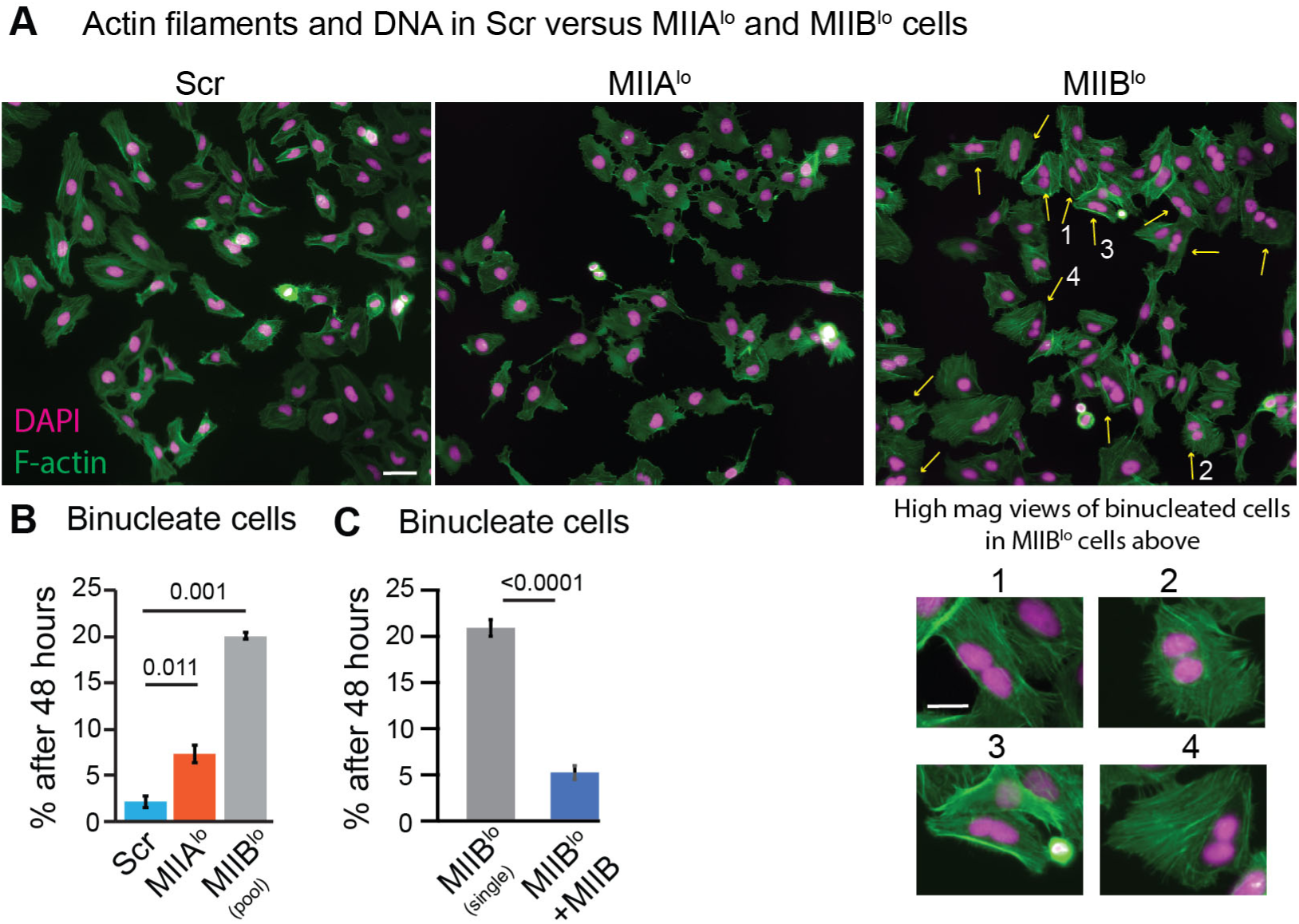
MIIB depletion leads to increase in binucleated cells. A) Representative fields of view of Scr, MIIA^lo^ and MIIB^lo^ cells 48 hours after re-plating showing F-actin (green) and nuclei (magenta). Inset for MIIB^lo^ shows examples of binucleated cells. B) Binucleate cells 48 hours post plating in Scr versus MIIA^lo^ (pooled siRNA) or MIIB^lo^ (pooled siRNA) cells. n= >1000 cells each over 3 independent experiments. C) Binucleate cells 48 hours post plating in MIIB^lo^ (single siRNA) versus MIIB^lo^ + MIIB cells. n= 760 MIIB^lo^ (single) and 584 MIIB^lo^ + MIIB cells over 3 independent experiments. Scale bar in (A)- 50 µm, inset-25 µm. Error bars represent standard error of the means.

### Loss of MIIB results in shape instabilities at the polar cortical network

The dramatic increase in binucleation indicated that something besides cleavage furrow ingression was altered in MIIB^lo^ cells. We analyzed MIIB^lo^ cells using high-resolution DIC imaging and observed MIIB^lo^ cells show dramatic shape instabilities during cytokinesis, in the form of large membrane blebs, while MIIA^lo^ cells showed reduced blebbing compared to Scr cells (Fig. 3A-C, Supplementary Movies 1-3). The large blebbing events in MIIB^lo^ cells caused spindle rocking behavior (Supplementary Movie 2), as has been previously upon destabilization of the polar cortex by inhibition of MII activity at one pole, causing cytokinetic failure (Sedzinski et al., 2011)

**Figure 3.**
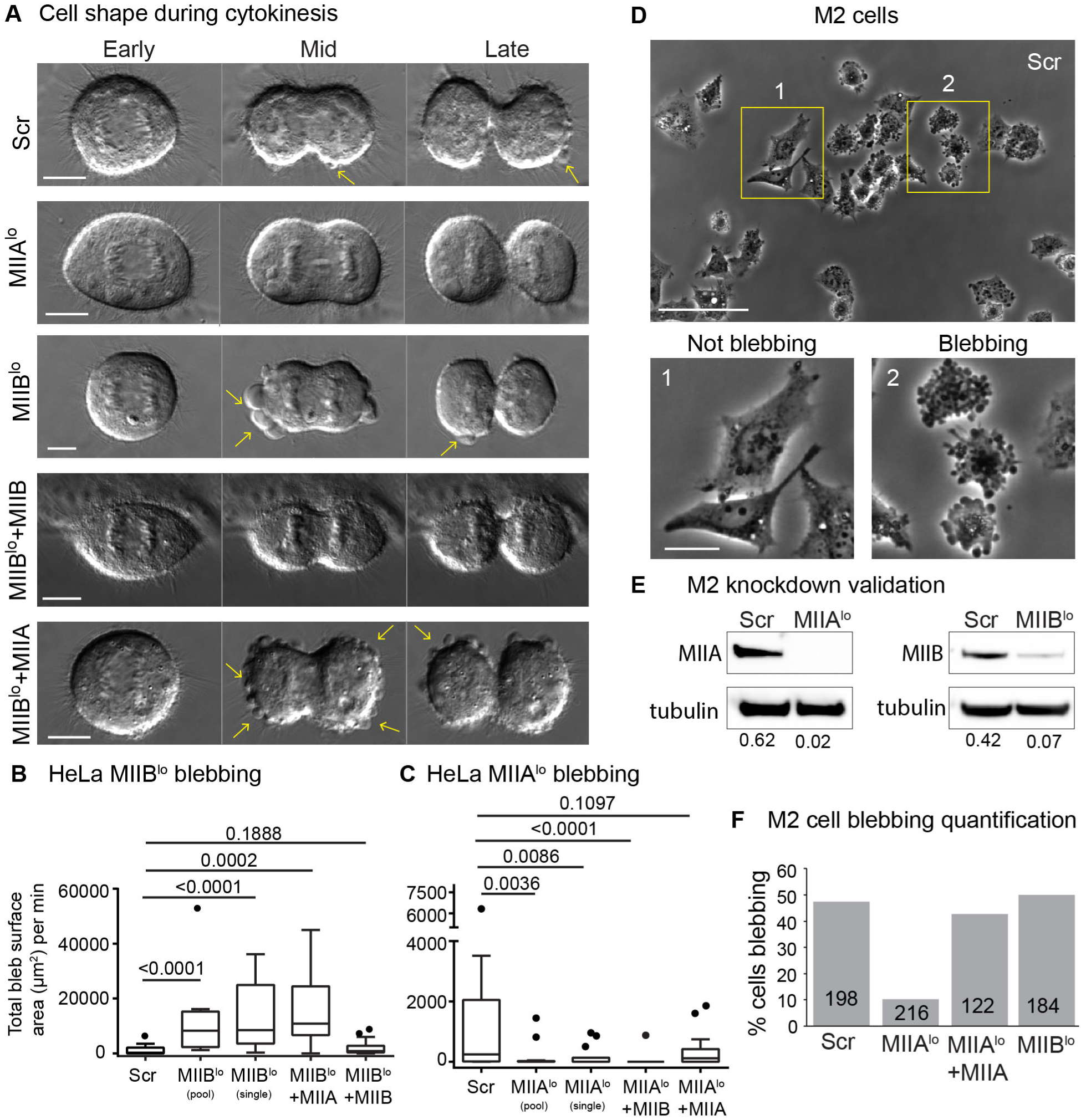
MIIB depletion leads to increased blebbing while MIIA depletion leads to decreased blebbing. A) Representative DIC time montages showing early, mid and late cytokinesis in Scr, MIIB^lo^, MIIA^lo^, MIIB^lo^+ MIIB-mEGFP and MIIB^lo^+ MIIA-mEGFP cells. Yellow arrows denote blebs. B-C) Tukey plots of degree of blebbing from time-lapse DIC measured as total bleb surface area per minute (B) n= 18 Scr cells, 15 MIIB^lo^ (pooled siRNA) cells, 16 MIIB^lo^ (single siRNA), 15 MIIB^lo^ +MIIB and 10 MIIB^lo^ +MIIA cells over 4 independent experiments. C) n=14 MIIA^lo^ (pooled siRNA), 14 MIIA^lo^ (single siRNA), 14 MIIA^lo^ +MIIA and 15 MIIA^lo^ +MIIB cells over 4 independent experiments. D) Representative field of control M2 5 hours post plating. Insets show examples of not blebbing (1) versus not blebbing (2) cells. E) Western blots validating knockdown MIIA and MIIB from M2 cells. Intensity of MII bands normalized to tubulin loading control shown below each band. F) Quantification of proportion of blebbing cells in Scr, MIIA^lo^, MIIB^lo^ or MIIA^lo^ + MIIA cells. The number of cells used in the quantification are stated inside each bar. Scale bars in (A)- 10 µm, (D)- 100 µm, inset-25 µm.

The increase in blebbing observed upon MIIB depletion was specific for MIIB, since expression of siRNA resistant MIIB-mEGFP at 119% of endogenous levels significantly reduced blebbing events in MIIB^lo^ cells (Fig. 3A, B). Overexpression of MIIA at 116% of endogenous levels in MIIB^lo^ cells did not reduce blebbing events (Fig. 3A, B; Supplementary Movie 4). This data suggested that this effect was specific to MIIB depletion. Conversely, to test whether reduction in blebbing observed in MIIA^lo^ cells was specific to MIIA depletion, we expressed siRNA resistant MIIA-mEGFP in MIIA^lo^ cells. We found no significant difference between Scr and MIIA^lo^ cells expressing MIIA-mEGFP (Fig. 3C). Overexpression of MIIB-mEGFP in MIIA^lo^ cells, however, did not increase blebbing (Fig. 3C, Supplementary Movie 5).

To further test if loss of MIIA is associated with reduced blebbing, we turned to filamin deficient M2 melanoma cells. M2 cells are a classical model for blebbing, since a large proportion of these cells constitutively blebs (Bovellan et al., 2014; Charras et al., 2006; Cunningham, 1995). We found ∼50% of Scr M2 cells blebbed. Knockdown of MIIA, but not MIIB, reduced the proportion of blebbing M2 cells (Fig. 3C-E, Supplementary Fig. 3). Expression of MIIA-mEGFP in MIIA^lo^ M2 cells increased the proportion of blebbing cells (Fig. 3E). Taken together, these data suggest loss of MIIA is correlated with lower blebbing, while loss of MIIB is correlated with higher blebbing.

The shape instabilities observed in MIIB^lo^ cells led us to further analyze cell shape upon depletion of the two paralogs. Specifically, we noted that MIIA^lo^ cells tended to elongate more than Scr cells during cytokinesis (Fig. 4A). We quantified this cell shape alteration by measuring the increase in pole-to-pole distance during cytokinesis relative to metaphase (Fig. 4B). Indeed, MIIA^lo^ cells showed a higher increase in pole-to-pole distance compared to Scr or MIIB^lo^ cells (Fig. 4C). To test if this increase in pole-to-pole elongation in MIIA^lo^ cells was specific to MIIA depletion, we expressed siRNA resistant MIIA-mEGFP in MIIA^lo^ cells. Expression of MIIA in MIIA^lo^ cells significantly reduced pole-to-pole elongation (Fig. 4D). On the other hand, overexpression of MIIB in MIIA^lo^ cells did not rescue this defect (Fig. 4D, Supplementary Movie 5). Taken together, our results show that depletion of either paralog leads to distinct alterations to cell shape during cytokinesis.

**Figure 4.**
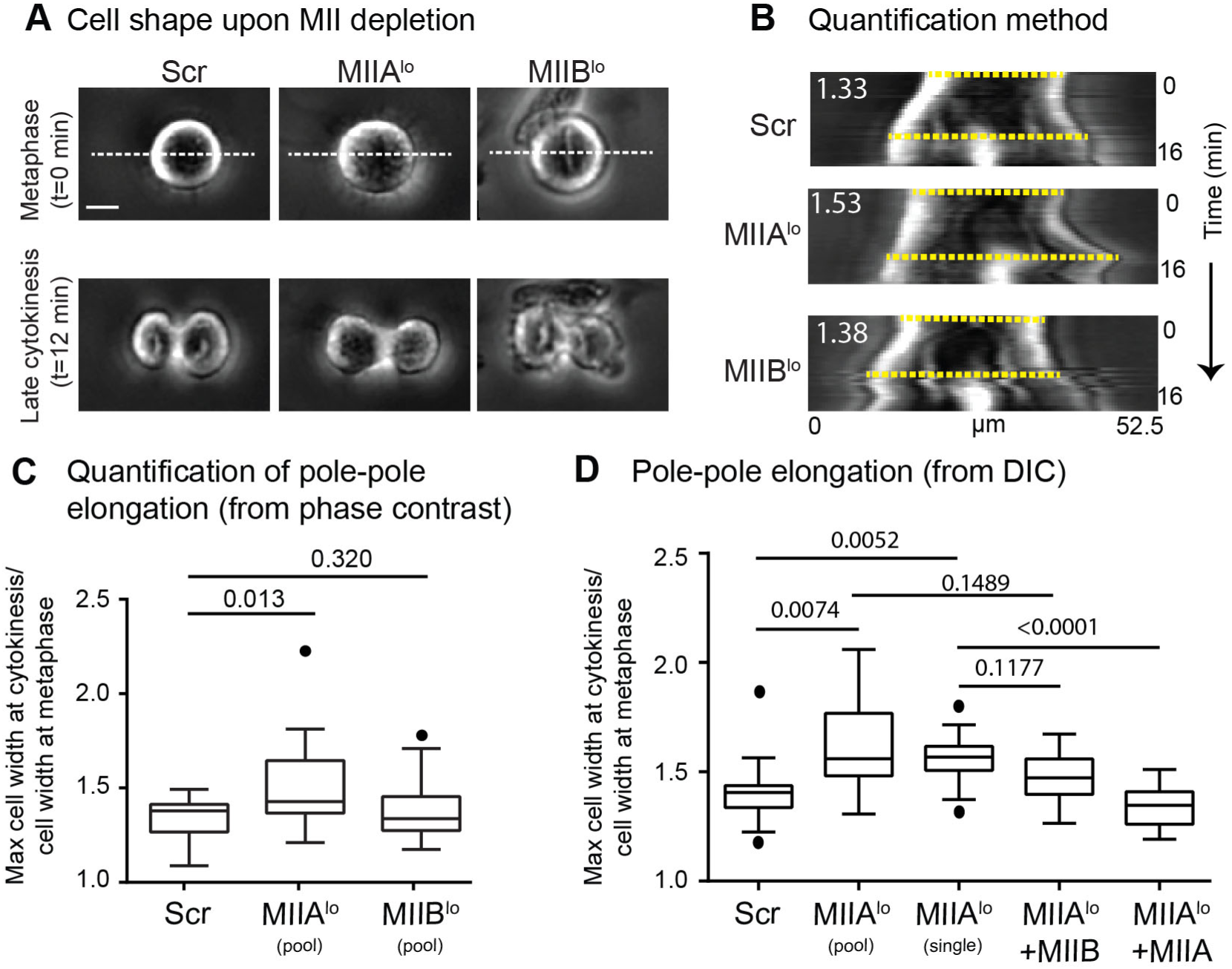
MIIA depletion leads to increased pole-to-pole elongation. A) Representative phase contrast images of Scr, MIIA^lo^ and MIIB^lo^ cells at metaphase and late cytokinesis. B) Kymographs were generated using dotted white lines in (A). Dotted yellow lines show the two pole-to-pole lengths used to measure the elongation ratio. See methods for detailed quantification method. C) Tukey plots of polar elongation measured using phase contrast. n=17 Scr, 20 MIIA^lo^ (pooled siRNA) and 18 MIIB^lo^ (pooled siRNA) cells over 3 independent experiments. D) Tukey plots of polar elongation measured independently using DIC. n= 15 Scr,12 MIIA^lo^ (pooled siRNA),13 MIIA^lo^ (single siRNA), 13 MIIA^lo^ +MIIA and 11 MIIA^lo^ +MIIB cells over 3 independent experiments. Scale bar- 10 µm. Solid circles represent outliers. p values stated over respective bars.

### MIIA and MIIB show similar spatiotemporal localization at the polar cortex during cytokinesis

To test whether the distinct phenotypes observed were driven by distinct localization of the two MII paralogs, we co-expressed MIIA mApple and MIIB mEmerald in HeLa cells and analyzed their spatial and temporal dynamics (Supplementary Movie 6). We noted MIIB was cortically enriched as early as metaphase, whereas MIIA was only weakly enriched at the metaphase cortex (Fig. 5A). After mitotic exit and prior to the onset of ingression (i.e., early cytokinesis), both MIIA and MIIB were enriched at the equatorial, and then at the polar cortex, colocalizing at the two networks. After the completion of furrow ingression (i.e., late cytokinesis), MIIA enrichment at the cortex was lost, while MIIB remained distinctly enriched at the equatorial cortex. To rule out over-expression artifacts and measure localization patterns in a quantitative manner, we characterized the endogenous localization of cortical MII populations and found that both MIIA and MIIB showed similar patterns of furrow and polar cortex enrichment as seen in our live cell data (Fig. 5B). These data show that while there were subtle differences in MIIA and MIIB localization, both paralogs localized to both the equatorial and polar cortex during furrow ingression. Other mechanisms must therefore account for the differences in phenotypes observed upon knockdown of MII paralogs, that is, the phenotypes observed are not due to only one paralog localizing to the polar cortex.

**Figure 5.**
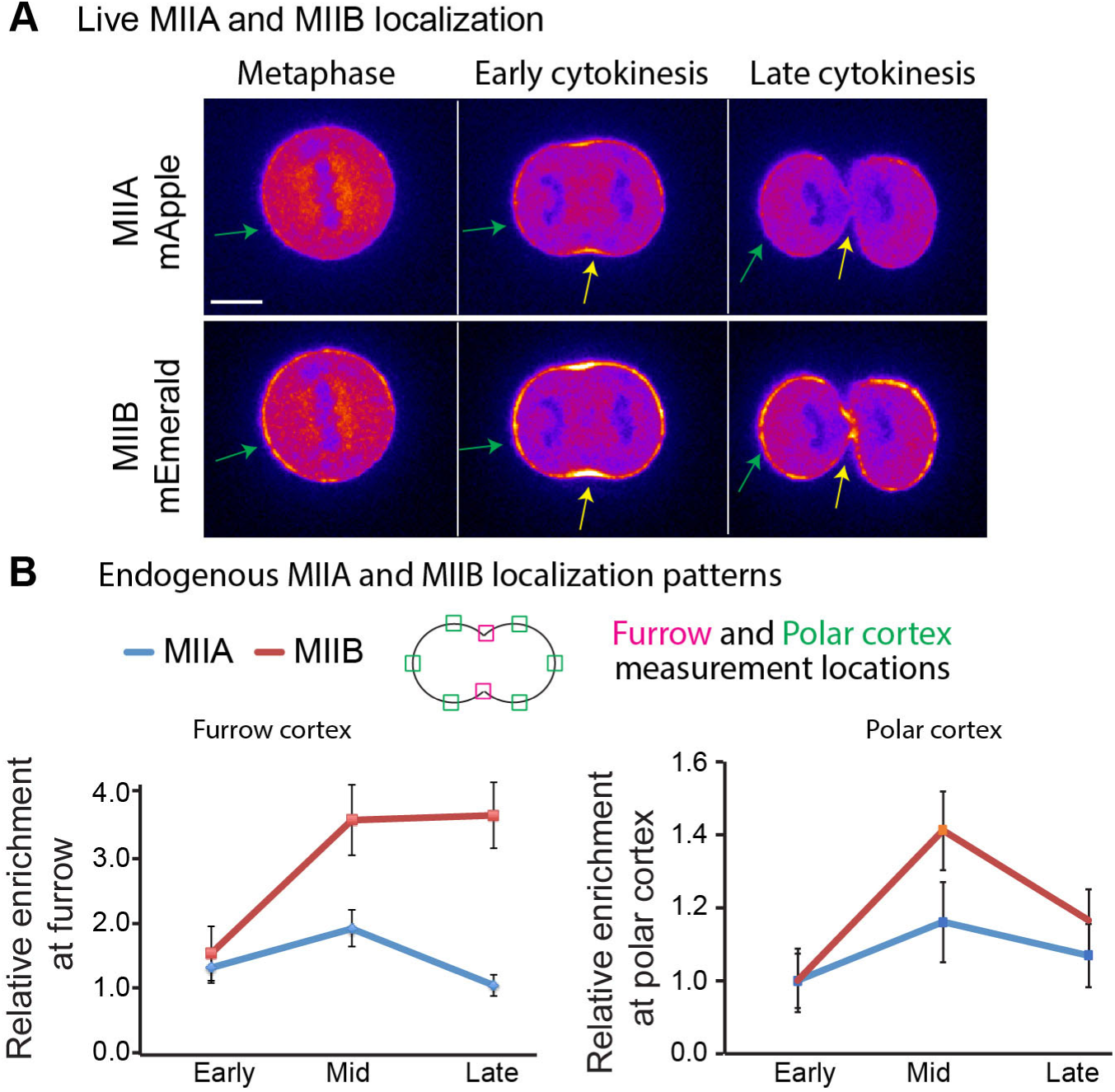
MIIA and MIIB show similar localization dynamics at the polar cortex. A) Live cell montage of HeLa cell co-expressing MIIA mApple and MIIB mEmerald. Green and yellow arrows highlight polar and equatorial cortex, respectively. B-C) Enrichment of endogenous MIIA and MIIB at the furrow (B) and the polar cortex (C) in control cells at mid, early and late cytokinesis. For MIIA: 42 cells measured over 3 independent experiments. For MIIB: 48 cells measured over 3 independent experiments. Representative images of control MIIA and MIIB endogenous localization can be found in Fig. 6A and B. Scale bar- 10 µm. Error bars represent standard error of the mean.

We hypothesized that the depletion of one paralog could lead to a change in the localization of the other. To test this, we localized endogenous MIIA and MIIB in Scr versus MIIB^lo^ and MIIA^lo^ cells, respectively. At the equatorial cortex, we noted only MIIB compensated for MIIA upon MIIA depletion (Supplementary Fig. 4). On the other hand, depletion of either paralog led to a compensatory increase in localization of the other at the polar cortex (Fig. 6A-C). We observed this compensatory behavior even at the metaphase cortex (Fig. 6D, E). Expression of MIIA in MIIA^lo^ cells at 60% of endogenous levels reduced MIIB localization at the cortex (Fig. 6F, G). Similarly, expression of MIIB in MIIB^lo^ cells at 123% of endogenous levels reduced MIIA localization at the cortex (Fig. 6F, G). Taken together, our data show that knockdown of either paralog leads to a change in the relative abundance of the two paralogs at the cortex.

**Figure 6.**
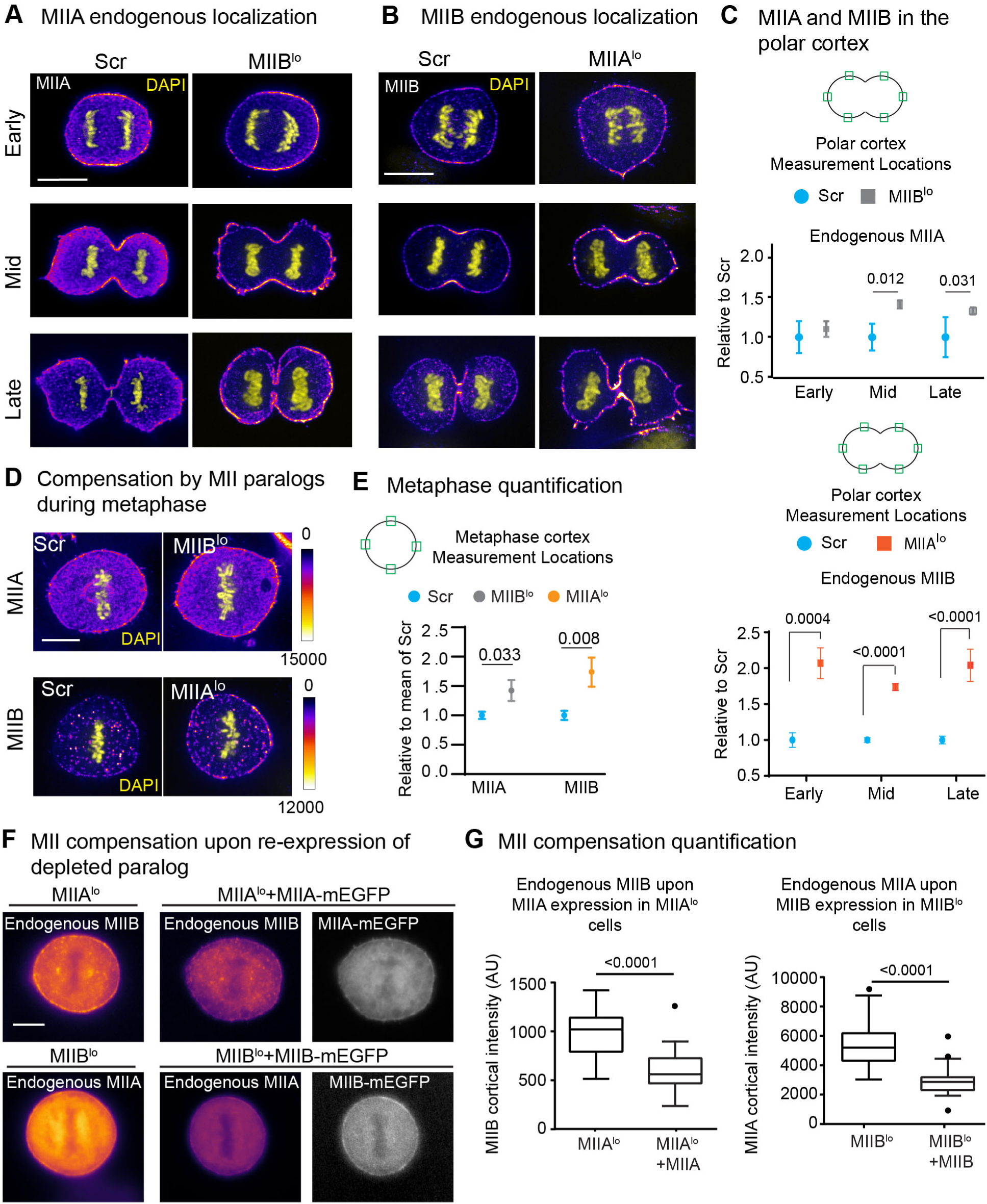
MII paralog compensation occurs at the metaphase and polar cortex upon MII knockdown. A-B) Early, mid and late cytokinesis cells stained for endogenous MIIA (A) or MIIB (B) (Fire LUT) comparing Scr versus MIIB^lo^ or MIIA^lo^ cells, respectively. All images compared were scaled similarly. C) Quantification of MIIA and MIIB at the polar cortex. Intensity was calculated as mean of green ROIs in cartoon insets. For MIIA: n=41 Scr and 48 MIIB^lo^ cells over 3 independent experiments. For MIIB: n= 46 Scr and 43 MIIA^lo^ cells over 3 independent experiments. D) Images showing MIIA or MIIB (Fire LUT) during metaphase in Scr versus MIIB^lo^ or MIIA^lo^ cells, respectively. MIIA: n=15 cells each for Scr and MIIB^lo^ over 3 independent experiments. MIIB: n= 13 Scr cells and 12 MIIA^lo^ cells over 3 independent experiments. E) Images showing MIIB localization in MIIA^lo^ versus MIIA^lo^ +MIIA-mEGFP cells and MIIB localization in MIIA^lo^ versus MIIA^lo^ +MIIA-mEGFP cells. F) Tukey plots for MIIB or MIIA cortical intensity. For endogenous MIIB, n= 28 MIIA^lo^ and 18 MIIA^lo^ +MIIA cells over 3 independent experiments. For endogenous MIIA, n= 34 MIIBlo and 25 MIIB^lo^ +MIIB cells over 3 independent experiments. Scale bars- 10 µm. Solid circles represent outliers. Error bars in (C) and (E) represent standard error of weighted mean.

### Depletion of MIIA results in reduced cortex tension and intracellular pressure

It has been previously reported that increased cortex tension results in increased bleb initiation and/or growth (Paluch et al., 2005; Tinevez et al., 2009). We therefore wondered whether the compensatory increase in the localization of one paralog upon depletion of the other altered cortex tension. Since MIIA depletion resulted in reduction in blebbing, we predicted that these cells would have lower cortex tension. Cortex tension has been classically measured using micropipette aspiration (Brugues et al., 2010; Kee and Robinson, 2013; Tinevez et al., 2009). We therefore performed micropipette aspiration on Scr and MIIA^lo^ cells (Figure 7A). We chose to perform these measurements during metaphase, since i) we find compensation in localization during metaphase ii) the metaphase cortex is uniform and is devoid of the fluctuations in MII localization that occur at the polar cortex (Supplementary Movie 6). iii) Polar cortex blebbing interferes in cortex tension measurements. We found a significant reduction (∼4-fold) in cortex tension in MIIA^lo^ versus Scr cells (Fig. 7B). This effect was specific to MIIA depletion since expression of MIIA-mEGFP at 72% of endogenous levels increased cortex tension to approximately control levels (Fig. 7B). As an additional control for these measurements, we treated Scr cells with 50 µM blebbistatin, an inhibitor of the ATPase activity of both MIIA and MIIB, and measured cortex tension. As previously reported, blebbistatin significantly reduced cortex tension (Tinevez et al., 2009) (Fig. 7B).

**Figure 7.**
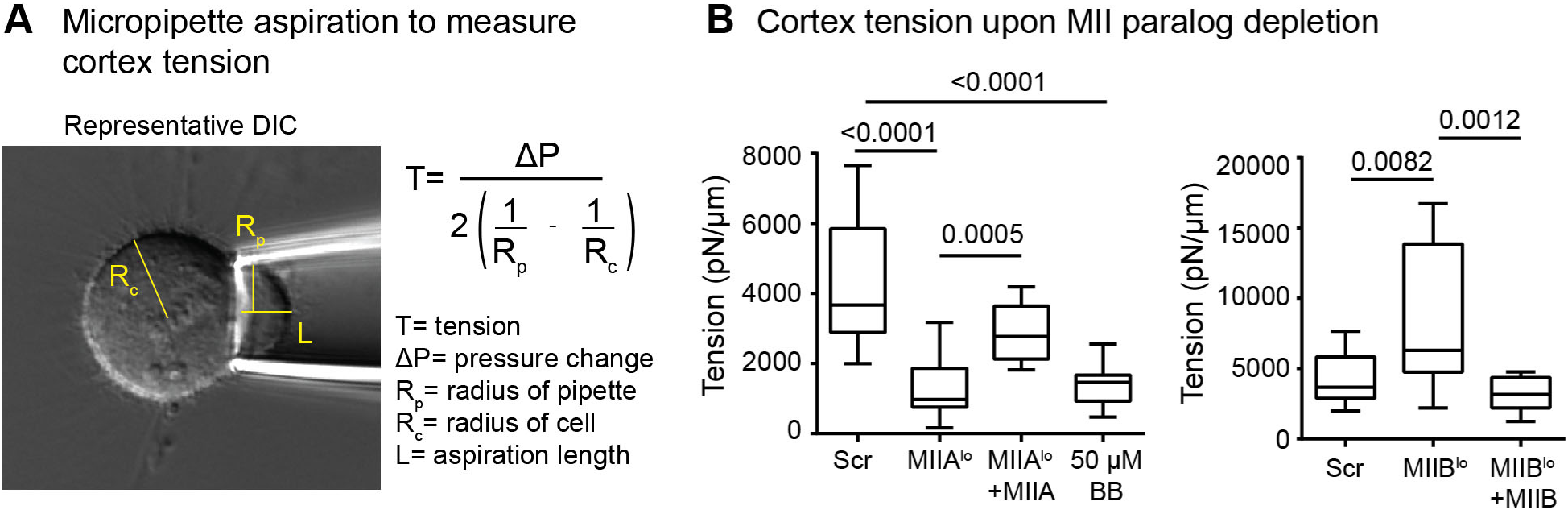
MIIA depletion leads to lower cortex tension while MIIB depletion leads to higher cortex tension. A) Representative image of micropipette aspiration of control cell during metaphase. Yellow lines represent radius of cell (R_c_), aspirated length (L) and radius of pipette (R_p_). Tension was calculated using the mathematical expression shown. B) Tukey plots of cortex tension upon MII paralog depletion. n= 16 Scr, 14 MIIA^lo^, 15 MIIB^lo^, 10 MIIA^lo^ +MIIA-mEGFP, 11 blebbistatin treated and 12 MIIB^lo^ +MIIB-mEGFP cells over 4 independent experiments each. P values stated over respective bars.

Conversely, we predicted that MIIB depletion should result in higher cortex tension as MIIB^lo^ cells show more blebbing. Indeed, MIIB^lo^ cells had ∼1.9 fold higher cortex tension compared to Scr cells (Fig. 7B). Expression of MIIB-mEGFP at 197% of endogenous levels in MIIB^lo^ cells resulted in a decrease in cortex tension comparable to Scr levels (Fig. 7B). Taken together, our results show MIIA depletion results in lower cortex tension, while MIIB depletion results in higher cortex tension, which correlated with the observed frequencies of bleb initiation we observed during cytokinesis (Fig. 3). Despite increased cortex tension in MIIB^lo^ cells, the cells did not bleb during metaphase. To test if increasing pressure could induce blebbing, we compressed MIIB^lo^ cells in metaphase with a microneedle. We observed MIIB^lo^ cells blebbed 80% of the time upon indentation, while Scr and MIIA^lo^ cells showed negligible blebbing (Supplementary Figure 5, Supplementary Movies 7-9).

We next wanted to directly interrogate the relative contributions of MIIA and MIIB to bleb growth. It has been previously reported that MII driven cortex tension promotes bleb growth by creating hydrostatic pressure (Tinevez et al., 2009). Blebs created by disruption of the cortex using localized laser ablation are thought to mimic the initiation of spontaneous blebs (Goudarzi et al., 2012; Tinevez et al., 2009). The size of the bleb created immediately following ablation is positively correlated with the instantaneous intracellular pressure. Ablation of the metaphase cortex of Scr cells resulted in the creation of a bleb, which subsequently retracted over a period of 45-90 seconds, as has been reported previously (Charras et al., 2006; Taneja and Burnette, 2019). Since MIIA^lo^ cells have lower cortex tension compared to Scr cells, we hypothesized that smaller blebs would be created in these cells. To test this, we created blebs using laser ablation at the metaphase cortex of MIIA^lo^ cells and measured the size of the bleb immediately following ablation (Fig. 8A). Small or no blebs were created immediately following ablation of the metaphase cortex in MIIA^lo^ cells (Fig. 8A-B). Interestingly, these blebs failed to retract and instead exhibited a period of slow growth over extended time periods of up to 4 minutes (Fig. 8A, Supplementary Fig. 6). This effect was specific to MIIA depletion, since expression of MIIA-mEGFP increased bleb sizes comparable to Scr cells, as well as restored bleb retraction (Fig. 8B). This role of MIIA in driving bleb retraction was consistent with our previous report (Taneja and Burnette, 2019).

**Figure 8.**
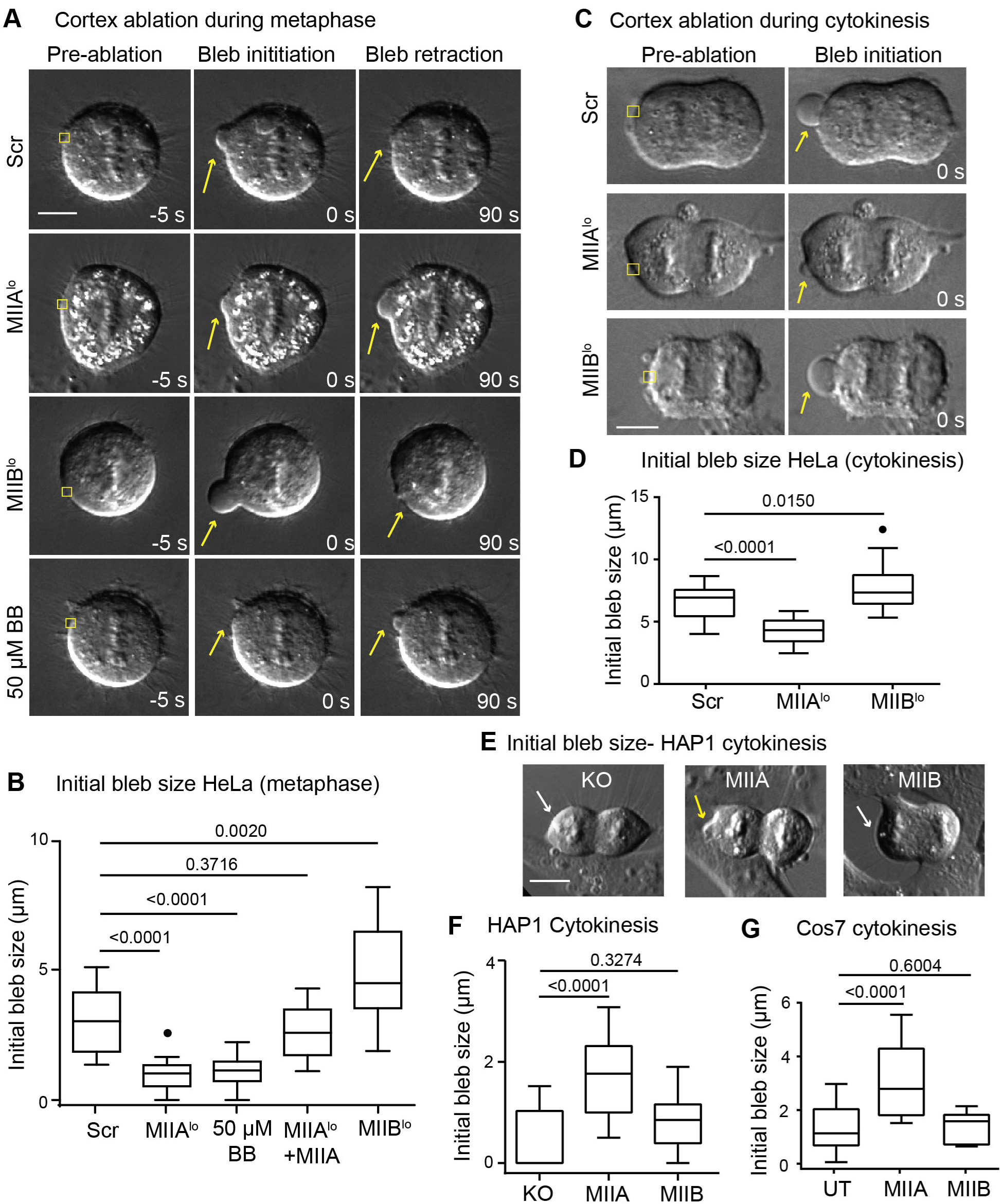
Loss of MIIA results in reduced tension and pressure while loss of MIIB results in increased tension and pressure. A) Representative images of Scr, MIIA^lo^, MIIB^lo^ and 50 µM blebbistatin treated HeLa before ablation, 0 seconds after ablation and 90 seconds after ablation of the metaphase cortex. Yellow squares depict ROI for ablation, yellow arrows depict blebs. B) Tukey plots showing initial bleb size during metaphase in HeLa cells. n= 17 Scr, 16 MIIA^lo^, 14 MIIA^lo^ +MIIA, 14 50 µM blebbistatin treated and 18 MIIB^lo^ cells over 3 independent experiments. C) Representative images of polar cortex ablation in Scr, MIIA^lo^ and MIIB^lo^ HeLa cells during cytokinesis. Yellow squares depict ablation ROI, and yellow arrows denote bleb created by ablation. D) Tukey plots of initial bleb size during cytokinesis in HeLa cells. n= 25 Scr, 15 MIIA^lo^ and 25 MIIB^lo^ cells over 3 independent experiments. E) Representative images of polar cortex ablation in HAP1 MIIA KO fibroblasts during cytokinesis. White arrows indicate no bleb was created while yellow arrows indicate creation of bleb following polar cortex ablation. F) Tukey plots of initial bleb size following ablation in HAP1 KO fibroblasts during cytokinesis. n= 12 untransfected MIIA-KO, 26 MIIA, and 11 MIIB expressing cells over 3 independent experiments. G) Tukey plots of initial bleb size following ablation during cytokinesis in Cos7 cells. n= 15 untransfected (UT), 15 MIIA and 11 MIIB expressing cells over 3 independent experiments. Scale bars- 10 µm. Solid circles represent outliers.

Conversely, we hypothesized larger blebs should be created upon depletion of MIIB due to increased cortex tension. Indeed, larger blebs were created in MIIB^lo^ cells upon laser ablation, and these blebs all successfully retracted (Fig. 8A-B, Supplementary Fig. 6). To test whether the pressure driving bleb growth was specifically generated by MII driven cortex tension, we performed ablation in cells treated with 50 µM blebbistatin. Indeed, blebbistatin treatment mimicked MIIA depletion, with small or no blebs created upon ablation, and exhibited slow bleb growth instead of retraction (Fig. 8A, B; Supplementary Fig. 6). Taken together, our results show MIIA driven cortex tension creates larger blebs upon ablation during metaphase.

We next proceeded to test whether MIIA drives growth of larger blebs at the polar cortex during cytokinesis. To that end, we performed laser ablation of the polar cortex during cytokinesis in Scr versus MIIA^lo^ and MIIB^lo^ cells (Fig. 8C). We observed that blebs created at the polar cortex of Scr cells resulted in blebs that were on average larger than those created during metaphase (Fig. 8D, compared to Fig. 8B), consistent with the idea that intracellular pressure increases following anaphase onset (Stewart et al., 2011). Depletion of MIIA resulted in the creation of smaller blebs that failed to retract, while MIIB depletion resulted in larger blebs compared to Scr cells (Fig. 8C-D).

We then wanted to test whether MIIA is sufficient to drive bleb growth. To that end, we performed laser ablation of the polar cortex of HAP1 MIIA KO cells. We observed that very small or no blebs were created in control KO cells (Fig. 8E, F). Expression of MIIA, but not MIIB, resulted in the formation of a bleb upon ablation (Fig. 8E, F). To further test if MIIA is sufficient to drive higher bleb growth, we turned to Cos7 cells, that do not express MIIA. Indeed, expression of MIIA, but not MIIB, increased bleb size upon ablation of the polar cortex during cytokinesis (Fig. 8G). Taken together, our results show that MIIA is both necessary and sufficient to drive the growth of larger blebs.

### MII motor domains dictate the relative contribution of MII paralogs

Previous studies have shown that the C-terminus of MII regulates paralog localization (Sandquist and Means, 2008; Vicente-Manzanares et al., 2008). The observed similarities in localization dynamics of MIIA and MIIB at the polar cortex (Fig. 5) suggest that the motor domains on the N-terminus might account for the distinct roles played by the two paralogs during cytokinesis. Therefore, a chimeric MII, bearing the motor domain of MIIB, and the coiled coil domain and tail piece of MIIA (MIIB/A), should be able to rescue blebbing in MIIB^lo^ cells by competing with endogenous MIIA motors at the polar cortex. As expected, this chimeric motor showed a localization pattern similar to MIIA, with weaker cortical enrichment (Fig. 9A), consistent with previous findings that the tail of MIIB determines its subcellular localization (Sandquist and Means, 2008; Vicente-Manzanares et al., 2008). Strikingly, expression of the MIIB/A chimera at levels comparable to full length MIIB (119% for MIIB/A versus 127% for MIIB) resulted in significant suppression of blebbing (Fig. 9B).

**Figure 9.**
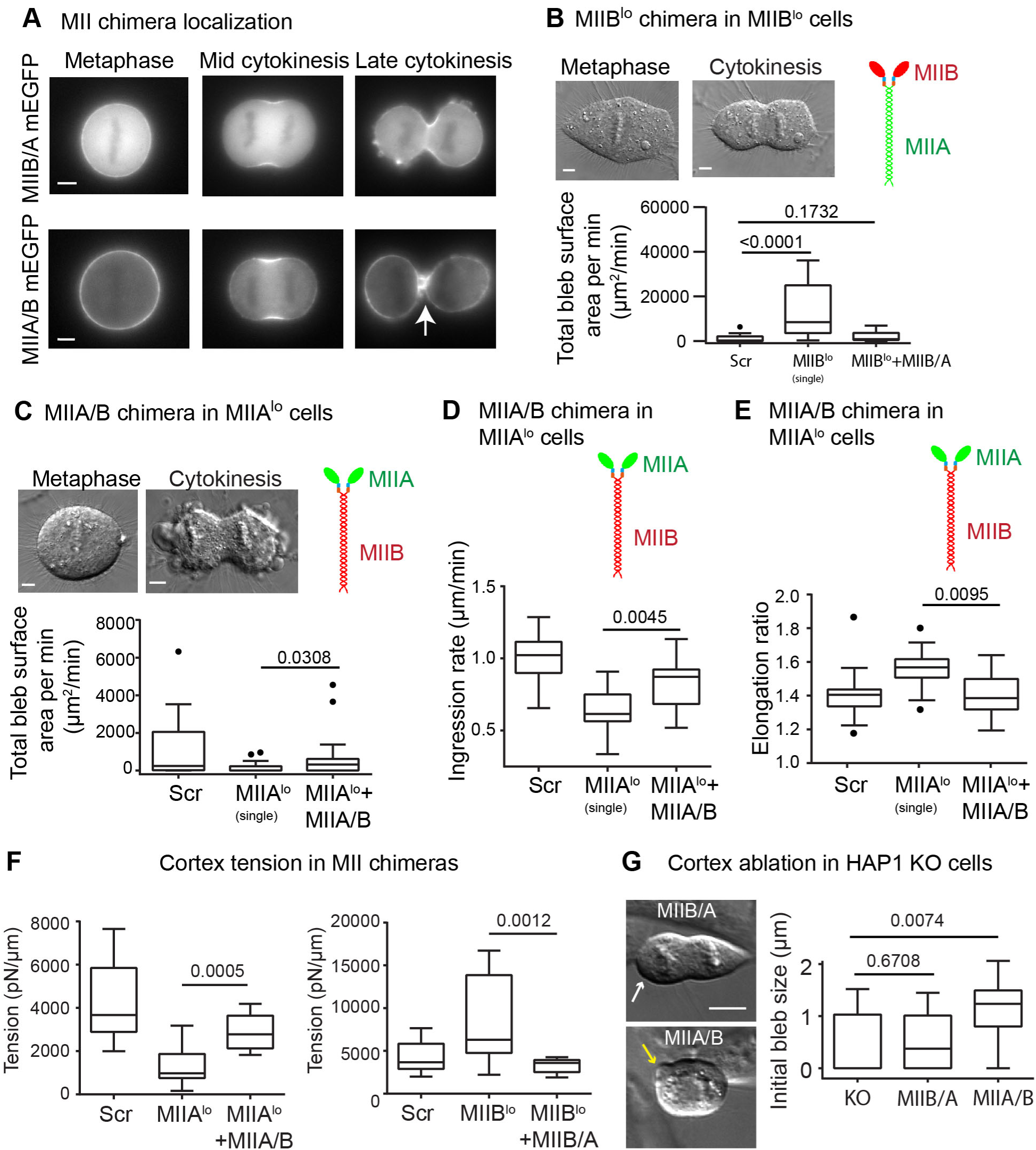
Motor domains determine the contribution of MII paralogs to polar cortex contractility. A) Localization of MIIB/A and MIIA/B chimera in MIIB^lo^ and MIIA^lo^ cells, respectively. B) Representative MIIB^lo^ cell expressing MIIB/A chimera at metaphase and cytokinesis. Tukey plots of blebbing events in MIIB/A expressing cells are compared to Scr and MIIB^lo^ (single siRNA). n=16 MIIB^lo^ +MIIB/A cells over 3 independent experiments. The Scr and MIIB^lo^ (single) datasets are the same as in Fig. 3B and are only shown for comparison. C) Representative MIIA^lo^ cell expressing MIIA/B chimera at metaphase and mid cytokinesis. n=16 cells MIIA^lo^ +MIIA/B cells over 3 independent experiments. The Scr and MIIA^lo^ (single) datasets are the same as Fig. 3C and are only shown for comparison. D) Tukey plots of cleavage furrow ingression rates from DIC time montages comparing MIIA^lo^ cells expressing MIIA/B chimera. n= 16 MIIA^lo^ +MIIA/B cells over 3 independent experiments. The Scr and MIIA^lo^ (single) datasets are the same as Fig. 1F and are only shown for comparison. E) Pole-to-pole elongation ratios for MIIA^lo^ cells expressing MIIA/B chimera. n=12 MIIA^lo^ +MIIA/B cells over 3 independent experiments. The Scr and MIIA^lo^ (single) datasets are the same as Fig. 4D and are only shown for comparison. F) Cortex tension in MIIB/A and MIIA/B chimeras using micropipette aspiration. n= 14 MIIA^lo^ +MIIA/B and 10 MIIB^lo^ +MIIB/A expressing cells over 4 independent experiments. The Scr, MIIA^lo^ and MIIB^lo^ datasets are the same as in Fig. 7B and are only shown for comparison. G) Initial bleb size following ablation in HAP1 KO cells. n=8 MIIB/A, 18 MIIA/B expressing cells over 3 independent experiments. Yellow and white arrow denote formation or absence of bleb, respectively. The control KO dataset is the same as Fig. 8F and is only shown for comparison. Scale bars in (A, G) and (B, C)- 10 µm and 5 µm, respectively. Solid circles represent outliers. P values stated over respective bars.

Conversely, MIIB is upregulated at the polar cortex in MIIA^lo^ cells (Fig. 6A-C); therefore, we hypothesized that increasing MIIA motors in the polar cortex should increase blebbing in MIIA^lo^ cells. To test this, we expressed a chimera bearing the motor domain of MIIA and the coiled coil domain and tailpiece of MIIB (MIIA/B) in MIIA^lo^ cells. This construct also faithfully mimicked MIIB localization, showing a clear cortical enrichment, as well as distinct equatorial localization during late cytokinesis (Fig. 9A). Interestingly, we observed a significant increase in blebbing upon expression of MIIA/B at levels comparable to full length MIIA (65% for MIIA/B versus 65% for MIIA) (Fig. 9C). Furthermore, this construct also rescued cleavage furrow ingression rates to control levels (Fig. 9D). In correlation with increased furrow ingression rates, cell shape alterations observed in MIIA^lo^ cells were also rescued by the MIIA/B chimera, with a decrease in pole-to-pole elongation (Fig. 9E). To test whether the suppression of the MIIA and MIIB depletion phenotypes correlated with changes in cortex tension, we performed micropipette aspiration on MIIA^lo^ expressing MIIA/B-mEGFP and MIIB^lo^ cells expressing MIIB/A-mEGFP. In line with our observations for ingression, blebbing and pole-to-pole elongation, we found MIIB/A expression in MIIB^lo^ cells significantly decreased cortex tension, while MIIA/B expression in MIIA^lo^ cells significantly increased cortex tension (Fig. 9F). To test whether these changes in cortex tension resulted in increased bleb growth, we expressed MIIA/B-mEGFP and MIIB/A mEGFP in HAP1 MIIA-KO cells and performed cortex ablation during cytokinesis. Expression of MIIA/B, but not MIIB/A significantly increased initial bleb size following ablation during cytokinesis (Fig. 9G). Taken together, these data suggest that the motor domain determines the paralog-specific contribution of the two motors at the polar and equatorial cortex.

## DISCUSSION

Recent biophysical studies have focused on actin architecture (Chugh et al., 2017) and cross-linker concentration (Ding et al., 2017) in the cortex as modulators of cortical mechanics during mitotic progression. MII is the major generator of tension in the cortex (Levayer and Lecuit, 2012; Salbreux et al., 2012), and while the role of MII contractility in regulating cortical tension has been intensively investigated in the context of bleb expansion (Tinevez et al., 2009) and mitotic cell rounding (Ramanathan et al., 2015; Stewart et al., 2011), these studies have treated contractility as a single parameter, lacking parameters such as differences in the properties of myosin paralogs. Our data shows that MIIA is responsible for both cleavage furrow ingression and the majority of cortex tension generation while MIIB is likely acting like an actin cross-linker stabilizing the polar cortex.

We found that MIIA depletion led to a reduction in the rate of cleavage furrow ingression (Figure 10). This reduction suggests that MIIA is the rate limiting motor driving ingression. However, it is important to note that MIIA depleted cells still ingressed their furrows and most completed cytokinesis, as we found only a minor increase in binucleated cells (Figure 2). On the other hand, MIIB depleted cells showed a marked increase in binucleation even though the rate of cleavage furrow ingression was not different from controls. This data supports previous suggestions that MIIB is important for the completion of abscission (Lordier et al., 2012) and a recent study has provided strong evidence that this is the case (Wang et al., 2019). In further support of this hypothesis, MIIB stayed enriched in the cleavage furrow at late stages of ingression whereas MIIA localization was reduced (Figure 5). Therefore, a failure of abscission could be the cause of the increase in binucleated MIIB depleted cells we observed (Figure 2). Another possible cause could be the rocking of the mitotic spindle that has been previously reported in cells with larger and more frequent membrane blebs (Sedzinski et al., 2011). As MIIB depletion correlated with such large blebbing events and spindle rocking (Supplementary Movie 2), it is possible that both sets of chromosomes could be positioned within a single daughter cell during cleavage furrow ingression.

**Figure 10.**
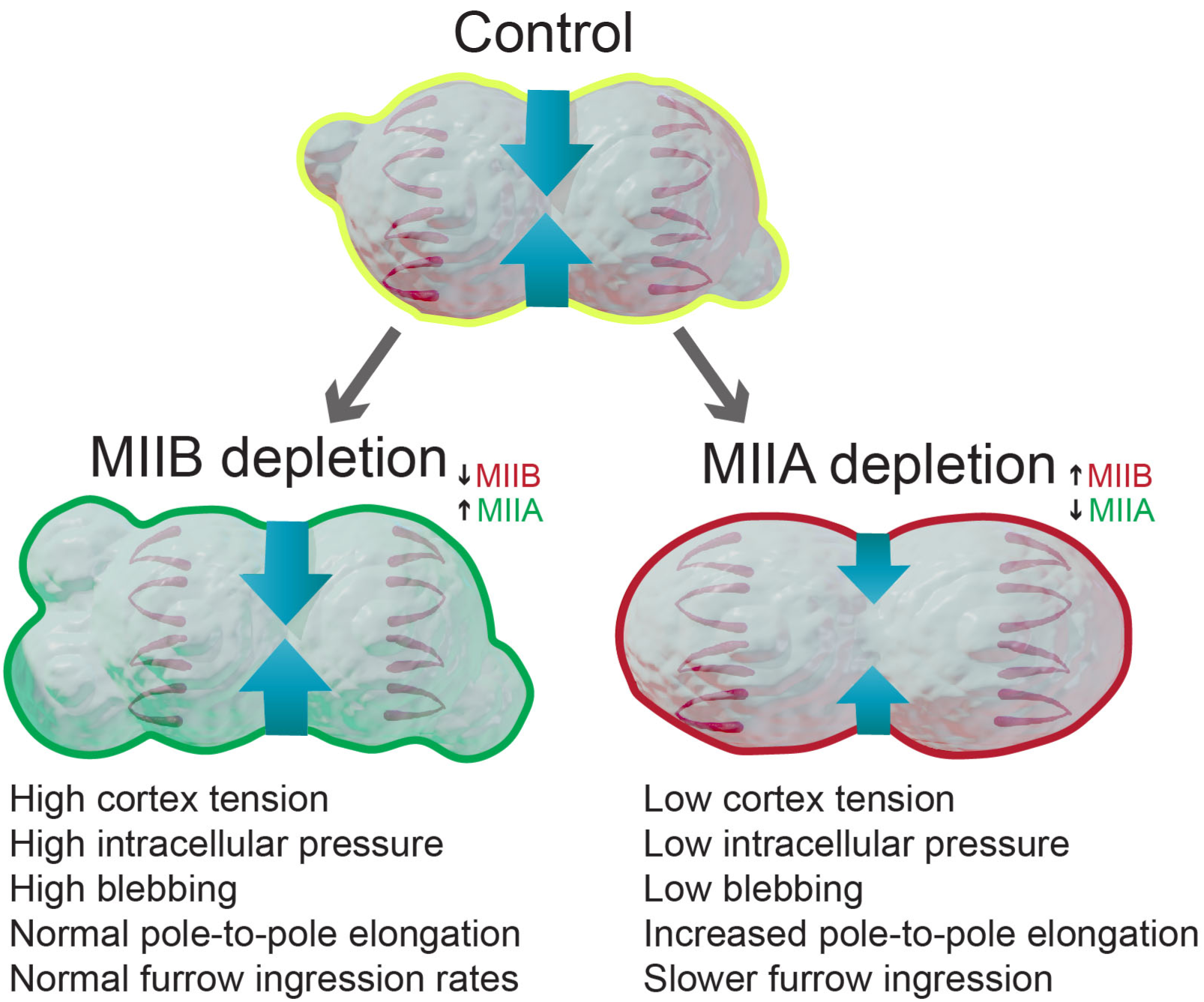
Working model. During cytokinesis, MIIA templates the formation of stacks of MIIA and MIIB at the cleavage furrow to drive ingression. Following onset of ingression, MIIA and MIIB both accumulate at the polar cortex, where their relative amounts determine their contribution to cortical tension and as a result, cortex stability. Under control conditions, they contribute equally, allowing normal amounts of blebbing to release excess pressure. Upon depletion of MIIB, MIIA localization increases at the polar cortex to increase cortex tension, in higher bleb nucleation, bleb growth that result in shape instability and cytokinetic failure. Upon depletion of MIIA, lower cortex tension correlates with slower cleavage furrow ingression, increased pole-to-pole elongation, in conjunction with loss of MII stack formation. At the polar cortex, MIIB compensates for MIIA and this results in lower cortex tension resulting in lower probability of bleb initiation and lower bleb growth. This results in decreased contractility and reduction in blebbing and increase in division failure.

The increased blebbing in MIIB depleted cells was our first clue that there were changes at the polar cortex. Our data shows MIIB depletion led to increased MIIA localization at the polar cortex, higher cortex tension and higher intracellular pressure. Previous experimental and theoretical studies have suggested the link between increased cortex tension and bleb initiation is driven by increased cortex breakage due to MII contractility (Paluch et al., 2005). At the molecular level, this may be driven by increased buckling of actin filaments (Murrell and Gardel, 2012). Increased cortex tension in MIIB depleted cells also resulted in increased bleb growth. Previous work has shown that increased cortex tension beyond a certain level drives the growth of very large blebs, which upon retraction, create large increases in pressure, which is sufficient to drive bleb initiation on the other side of the cell (Sedzinski et al., 2011). This type of positive feedback may be driving cell shape oscillations we observe in MIIB depleted cells. This phenotype is in contrast with the very different shape change we observed upon MIIA depletion.

A reduction in cortex tension in MIIA depleted cells correlated with an increase in cell elongation during cytokinesis (Figure 10). We speculate there are two possible mechanisms that could account for this shape change. First, it is possible that decreased furrow contractility— as suggested by reduced rates of ingression— could lead to the daughter cells “pulling” apart through a substrate adhesion dependent mechanism. Indeed, a recent study has proposed this mechanism in cells lacking a contractile ring due to depletion of Ect2 (Dix et al., 2018). Second, reduced cortex tension at the polar cortex could allow for an elongation of the mitotic spindle during anaphase. These and other potential mechanisms should be the focus of future studies.

Our data shows that the motor domain of each MII paralog determines its contribution to cortex tension, with lower duty ratios and higher ATPase activity of MIIA associated with higher cortex tension. Depletion of either paralog leads to a compensatory increase in localization at the polar cortex by the other. MIIA depletion leads to increased MIIB localization at the polar cortex, resulting in reduced polar MIIA contractility. This could lead to increased cortical stability, as shown by lower cortex tension and less bleb initiation and growth. MIIB depletion, on the other hand, leads to increased MIIA localization at the polar cortex, resulting in increased MIIA polar contractility and higher cortex tension. The precise mechanism by which this compensation tunes cortex tension at the molecular level is still unclear.

On the molecular level, it has been well established that MIIA and MIIB co-assemble into hetero-filaments by interaction of their coiled-coil domains (i.e., a bipolar MII filament containing both paralogs) (Beach et al., 2014; Shutova et al., 2014). A recent *in vitro* study has shown the relative number of MIIA and MIIB filaments within hetero-filaments of the two motors can modulate the ability of a hetero-filament to processively walk along actin filaments (Melli et al., 2018). Our data suggests that the relative levels of MIIA and MIIB at the cortex can also modulate cortex tension. Reducing MIIA resulted in lower cortex tension, while reducing MIIB resulted in higher cortex tension. Therefore, it is possible that fine-tuning of cortex tension at the molecular level occurs by varying amounts of MII motors within co-filaments. Alternatively, MIIA and MIIB could each form discrete filaments and occupy discrete sites on the polar cortex. Future studies using immuno-gold electron microscopy will be needed to distinguish between these two possibilities.

Depleting MIIA resulted in lower cortex tension, despite an increase in MIIB localization at the cortex. We were not surprised that MIIB did not play a larger role in generating cortex tension. MIIB has a higher duty ratio than MIIA (Kovacs et al., 2003; Rosenfeld et al., 2003; Wang et al., 2003). That is, MIIB binds actin in the force generating state for a larger proportion of its mechano-chemical cycle. Therefore, the canonical view of MIIB has been that it is adapted to maintain tension rather than generate tension. This view has been supported by theoretical studies on cortex tension generation mechanisms that have also implicated MIIB’s role as a cross-linker at the cortex since it is a slower motor (Stam et al., 2015).

The vast majority of non-muscle contractile systems observed in nature are cortical (i.e., associated with the plasma membrane), and the modulation of these contractile systems can drive diverse processes during development and disease progression (Levayer and Lecuit, 2012; Salbreux et al., 2012). Bleb based migration is a major mode of motility employed during early embryogenesis (Blaser et al., 2006; Holtfreter, 1943; Kageyama, 1977; Trinkaus, 1973) as well as cancer metastasis (Friedl and Wolf, 2003; Sahai and Marshall, 2003). MIIA upregulation is known to associate with metastasis in many solid tumors (Derycke et al., 2011; Maeda et al., 2008). Our working model would predict that a high MIIA to MIIB ratio would predispose these cells to use bleb-based migration. Furthermore, MII paralog expression is also very dynamic during hESC differentiation, correlating with different germ layer lineages (Sato et al., 2003). These changes in paralog expression may have important roles in regulating cortical mechanics during both cell migration and division across cell types from various non-muscle cellular systems.

Roles for MII paralogs and their regulatory kinases have been proposed in multiple other contexts, such as stress fiber formation in migrating cells (Beach et al., 2017), junction formation in epithelial cells (Smutny et al., 2010), growth cone advance in neurons (Brown and Bridgman, 2003), ESC differentiation (Rosowski et al., 2015; Sato et al., 2003) and pro-platelet formation in mice (Bluteau et al., 2012; Lordier et al., 2012). In some instances, such as cell migration and epithelial junction formation, the differential roles are exerted by regulation of localization. In epithelial cells, Rho/ROCK recruits MIIA while Rap1 recruits MIIB (Smutny et al., 2010). In other instances, such as megakaryocyte maturation, MIIB is transcriptionally downregulated to promote cytokinetic failure (Lordier et al., 2012). During proplatelet formation, the duty ratio of MIIA is known to be essential for platelet release (Badirou et al., 2015). Here we delineate the differential contributions of MIIA and MIIB at the polar cortex during cytokinesis, and our results are likely to have implications for cortical mechanics in these systems.

Finally, it is interesting to note that organisms such as yeast, worms, and flies only contain one MII paralog, leading to the question of how contractility is tuned in these organisms. One potential mechanism may be increased reliance on adhesion dependent cytokinesis. Indeed, organisms such as *Dictyostelium* and adhesive cell lines such as normal rat kidney (NRK) cells can divide without MII activity (Kanada et al., 2005; Neujahr et al., 1997). Another mechanism recently proposed could involve regulating the amount of each paralog that can assemble into filaments, for instance through PKC activity (Schiffhauer et al., 2019). Finally, oscillations of upstream regulators of MII, which appear to be evolutionarily conserved, and has been observed from flies to mammals (Bement et al., 2015; Machacek et al., 2009; Martin et al., 2009), could regulate the level of MII activation. The mechanism we propose here adds an additional layer of control and tunability, likely acquired later in evolution, when MII paralogs diverged.

## ACKNOWLEDGEMENTS

We thank David Miller for providing glass micro-capillaries for the indentation assay and Miguel Vincente-Manzanares for providing MII chimeras. We thank the Nikon Center of Excellence, Vanderbilt University for providing access to the Nikon Spinning Disk microscope and technical support. We thank Kristopher Burkewitz for generously letting us borrow his InjectMan for our micro-pipette aspiration experiments after our TransferMan malfunctioned. This work was funded by a MIRA from NIGMS (R35 GM125028-01) to D.T.B, an American Heart Association Predoctoral Fellowship (18PRE33960551) to N.T. and a MIRA (R35-HL135790) to W.D.M. The authors declare no competing financial interest.

## AUTHOR CONTRIBUTION

D.T.B and N.T. designed experiments. N.T., S.B. and A.M.F performed experiments. N.T, M.R.B, A.M.F and J.A.C analyzed the data. M.R.B and W.D.M helped in the interpretation of cell indentation experiments. R.O. helped in experiment design. V.G provided access to reagents (human H9 ESC cell line). N. T and D.T.B wrote the manuscript. All authors read and commented on the manuscript.

## MATERIALS AND METHODS

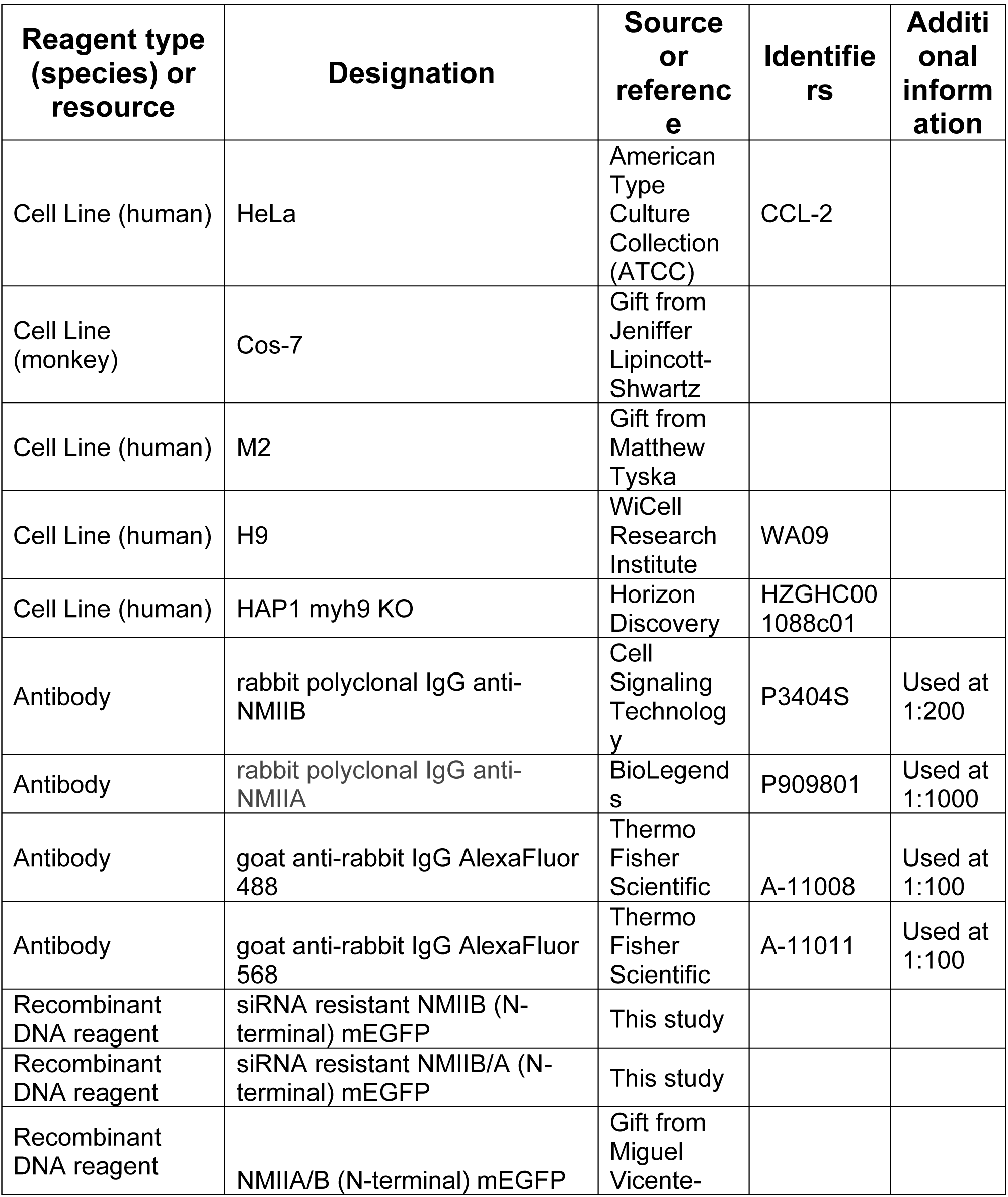
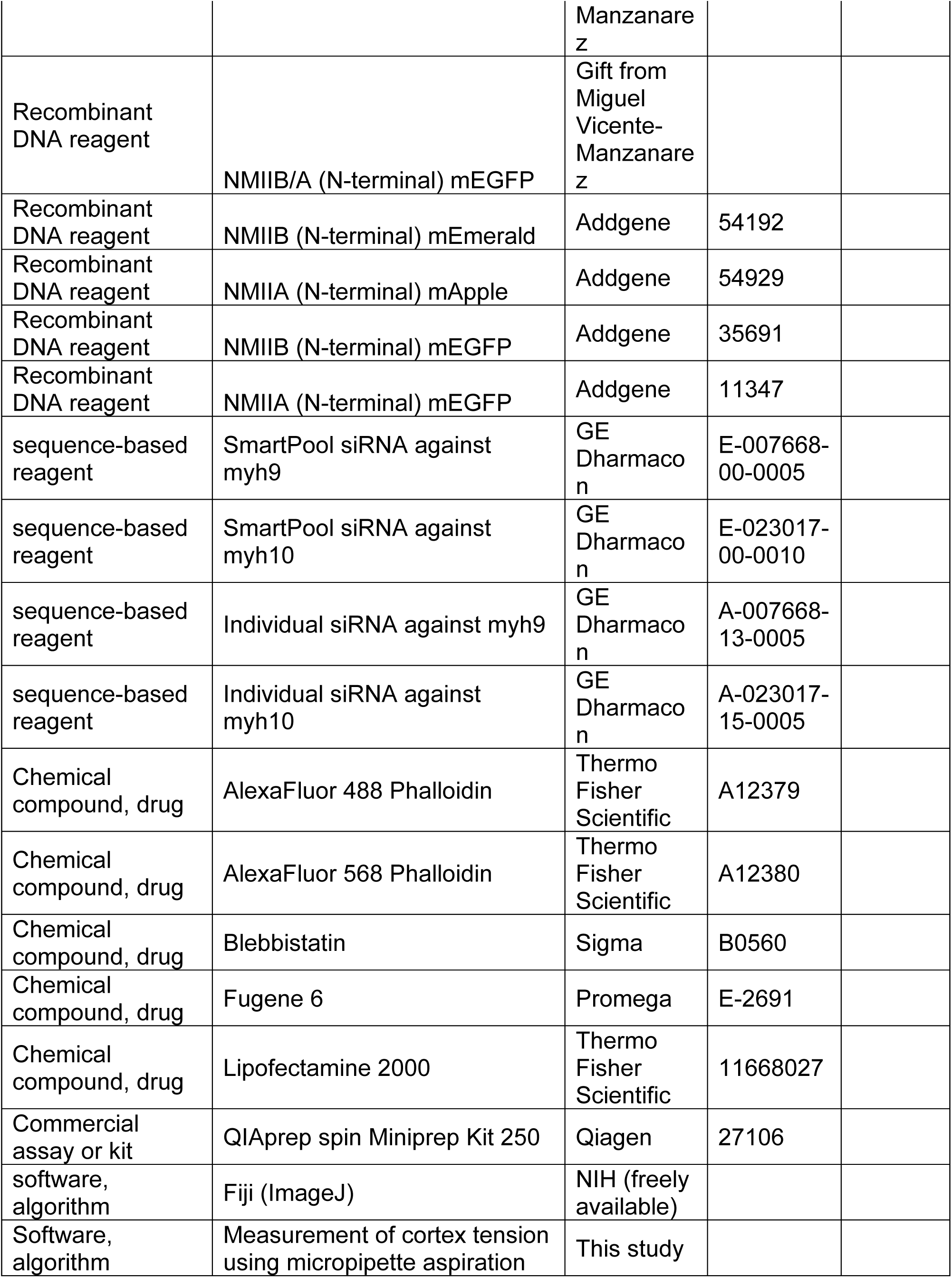
Key Resources Table

### Cell lines and growth conditions

HeLa (ATCC, CCL-2) and Cos7 cells (ATCC, CRL-1651) were cultured in growth media comprised of DMEM (Mediatech, Inc., Manassas, VA, #10-013-CV) containing 4.5 g/L L-glutamine, L-glucose, sodium pyruvate and supplemented with 10% fetal bovine serum (Sigma-Aldrich, St. Louis, MO, #F2442). HAP1 *myh9* (MIIA) KO and parental cells were purchased from Horizon discovery as previously described (Fenix *et al*., 2016), and cultured in IMDM medium supplemented with 10% fetal bovine serum. M2 melanoma cells were cultured Minimal Essential Medium supplemented with Earle’s salts, 10 mM HEPES and 10% fetal bovine serum. Growth substrates were prepared by coating #1.5 glass coverslips (In Vitro Scientific, #D35C4-20-1.5N or #D35-20-1.5N) with 10 μg/mL FN (Corning, Corning, NY, #354008) in PBS (Mediatech, Inc., #46-013-CM) at 37 °C for 1 hour.

Human embryonic stem cell lines H9 (WA09) were obtained from WiCell Research Institute (Madison, WI). Cells were seeded as undifferentiated colonies on plates coated with Matrigel (Corning, #354277), maintained at 37°C and 5% CO_2_, and fed daily with mTeSR (Stem Cell Technologies #05850). Cells were passaged once they reached 80% confluency. For immunofluorescence, cells were plated on Matrigel-coated 35 mm glass bottom culture dishes (MatTek #P35G-0-140C). Cells were plated on a growth substrate and then experiments were performed the next day.

For protein expression, cells were transiently transfected using Fugene 6 (Promega, Madison, WI, #E2691) as per the manufacturer’s instructions overnight in a 25-cm^2^ cell culture flask (Genessee Scientific Corporation, San Diego, CA, #25-207) before plating on a growth substrate.

Alexa Fluor-488 phalloidin (#A12379), Alexa Fluor-568 phalloidin (#A12380) and Alexa Fluor 488-goat anti-rabbit (#A11034) were purchased from Life Technologies (Grand Island, NY). Rabbit anti-myosin IIA (#909801) was purchased from BioLegends (San Diego, CA). Rabbit anti-myosin IIB (#8824S) was purchased from Cell Signaling Technology (Danvers, MA).

### Cell Line Authentication

The HeLa cell line used in this study was a gift of Dr. DA Weitz (Harvard University), and the M2 melanoma cells were a gift from Dr. Matthew Tyska. The Burnette lab had these lines authenticated by Promega and ATCC using their ‘Cell Line Authentication Service’ in 2015 and 2019, respectively. The methods and test results received from Promega and ATCC are as follows:

“Methodology: Seventeen short tandem repeat (STR) loci plus the gender determining locus, Amelogenin, were amplified using the commercially available PowerPlex 18D Kit from Promega. The cell line sample was processed using the ABI Prism 3500xl Genetic Analyzer. Data were analyzed using GeneMapper ID-X v1.2 software (Applied Biosystems). Appropriate positive and negative controls were run and confirmed for each sample submitted.’

“Data Interpretation: Cell lines were authenticated using Short Tandem Repeat (STR) analysis as described in 2012 in ANSI Standard (ASN-0002) Authentication of Human Cell Lines: Standardization of STR Profiling by the ATCC Standards Development Organization (SDO)’

#### HeLa- CCL-2 results

“Test Results: The submitted profile is an exact match for the following ATCC human cell line(s) in the ATCC STR database (eight core loci plus Amelogenin): CCL-2 (HeLa)’

#### M2 results

“Test Results: Submitted sample, STRA12409 (M2 melanoma), is an exact match to ATCC cell line CRL-2500 (A7). When compared to the reference profile the submitted profile shows and extra #10 allele at the TPOX locus. The cell line, (M2), has been discontinued by ATCC.”

Mycoplasma Monitoring: All cell lines were checked for potential mycoplasma infection using DAPI throughout the course of this study.

### Plasmids

MIIA mEmerald (Addgene, Cambridge, MA, #54190), MIIA mApple (Addgene, #54929) and MIIB mEmerald (Addgene, #54192) were gifts from Michael Davidson. MIIA mEGFP (Addgene, #11347) was a gift from Robert Adelstein. MIIB mEGFP (Addgene, #35691) was a gift from Venkaiah Betapudi. The MIIB/A and MIIA/B chimeras were generously provided by Miguel Vincente Manzanares (Universidad Autonoma de Madrid, Spain).

siRNA resistant MIIB and MIIB/A were generated using a modified site directed mutagenesis protocol as described previously (Liu and Naismith, 2008). Three to four codons were modified to ensure siRNA resistance. The MIIB- and MIIB/A-mEGFP constructs were mutated using the following primer sequences-Forward-5’-GGAAGACCCCGAGAGGTATCTCTTTGTGGACAGGGCTGT-3’, Reverse-5’-CTCTCGGGGTCTTCCAGTCCAGTTCTCTGCGCCAT-3’. Since the individual siRNA against MIIA was targeted to 3’-UTR, the MIIA-mEGFP and MIIA/B-mEGFP plasmids were not mutated.

### Phase and DIC imaging

Phase and DIC imaging was performed on a Nikon (Melville, NY) Eclipse Ti-E inverted microscope equipped with a Nikon 1.45 NA 100X Oil DIC, 0.95 NA 40X Air DIC and a 0.4 NA 20X Air Phase objective. Samples were maintained at 37°C with 5% CO_2_ using a Tokai Hit Stage Incubator (Shizuoka-ken, Japan).

### Structured Illumination Microscopy (SIM)

SIM imaging was performed on a GE (Pittsburgh, PA) DeltaVision OMX microscope equipped with a 1.42 NA 60X objective lens and a sCMOS camera.

### Live and Fixed Fluorescence imaging and Cortical Ablation

Live imaging and cortical ablation were performed on a Nikon Eclipse Ti-E inverted microscope equipped with a Yokogawa CSU-X1 spinning disk head, 1.4 NA 60X oil objective, Andor DU-897 EMCCD and a dedicated 100 mW 405 diode ablation laser, generously provided by the Nikon Centre of Excellence at Vanderbilt University. The instrument was controlled using Nikon Elements AR software. For ablation, a 1.4 µm x 1.4 µm ROI was used for all experiments. A DIC and/or fluorescence image was acquired before ablation, followed by ablation using a miniscanner. A pixel dwell time of 500 µs, 50% laser power was used for a duration of 1 second, followed by acquiring DIC or fluorescence images at 2 second intervals. Samples were maintained at 37°C with 5% CO_2_ using Tokai Hit Stage Incubator.

To image MIIA and MIIB in fixed ESC colonies, large image stitching was performed using the 60X objective in Elements software. Z-sections were acquired at 1 µm intervals for the entire stitch and maximum projections were displayed and used for generating line scans. To image endogenous MIIA and MIIB in fixed HeLa cells, single slices through the middle of the cell were acquired using the 60X objective.

### Knockdown experiments

Smart Pool Accell siRNA against MIIA (myh9 gene, #E- 007668, #1- CCGUUGACUCAGUAUAGUU, #2- UCCACAUCUUCUAUUAUCU, #3- GUGUGGUCAUCAAUCCUUA, #4- CUUAUGAGCUCCAAGGAUG) and MIIB (myh10 gene, #E- 023017, #1- GGACUAAUCUAUACUUAUU, #2- UGUCAAUGCUUAAAGUAGU, #3- CGAGGAUCCAGAGAGGUAU, #4- CCAAUUUACUCUGAGAAUA) were purchased from GE Dharmacon (Lafayette, CO). To perform rescue experiments, the following individual siRNAs were used-myh9 – CCGUUGACUCAGUAUAGUU (#A-007668-13-0005) and myh10-CGAGGAUCCAGAGGUAU (#A-023015-15-0005). Knockdown experiments were performed in 24-well plates using Lipofectamine 2000 (Life Technologies, #1690146) as per instructions provided by the manufacturer. Knockdown was performed for 72 hours, after which cells were either plated on the growth substrate for imaging or lysed for western blot experiments.

### Western Blotting

Gel samples were prepared by mixing cell lysates with LDS sample buffer (Life Technologies, #NP0007) and Sample Reducing Buffer (Life Technologies, #NP00009) and boiled at 95°C for 5 minutes. Samples were resolved on Bolt 4-12% gradient Bis-Tris gels (Life Technologies, #NW04120BOX). Protein bands were blotted onto a nylon membrane (Millipore). Blots were blocked using 5% NFDM (Research Products International Corp, Mt. Prospect, IL, #33368) in Tris Buffered Saline with Tween-20 (TBST). Antibody incubations were also performed in 5% NFDM in TBST. Blots were developed using the Immobilon Chemiluminescence Kit (Millipore, #WBKLS0500).

### Cell indentation assay

A precisely controlled Transfer Man 4R micromanipulator (Eppendorf, Hamburg, Germany) was magnetically attached to the optical table. Indentation pipettes were prepared from borosilicate microcapillary tubes pulled using an automated pipette puller. Metaphase cells were identified using high magnification DIC and their height was approximated. The micropipette was then slowly lowered to gently rest on top of the cell. Upon starting image acquisition, the pipette was lowered continuously for 2 seconds, using the extra-fine movement setting, to compress the cell to half its height. Images were recorded for a minimum of 90 s prior to gently raising the pipette back up. The assay was performed in 35 mm glass bottom dishes with a 20 mm coverslip to allow for the optimal angle and movement of the microcapillary.

### Micropipette Aspiration

Microneedles (World Precision Instruments, 0.75 mm inner diameter and 1.0 mm outer diameter) were pulled to a centimeter-long taper using a Narishige PC-100 needle puller. The following parameters were used-55% maximum current and one light weight (mass = 23.5g). Using a Narishige MF-900 microforge, needles were then cut to a diameter ranging between 10-15 µm. The end of the needle was then fire polished by bringing it close to a heated glass bead to obtain a smooth edge that will successfully attach to the aspirated cell without disrupting the membrane. The microforge was then used to bend the tip of the micropipette needle to a 30-degree angle by positioning the needle vertically at a 0.1 mm distance from the heated glass bead.

To perform micropipette aspiration, the needles were mounted on an Eppendorf Transfer Man 4R micromanipulator that was magnetically attached to the optical table. Cells were visualized using DIC (Plan Apo 0.95 NA, 40X Nikon DIC air objective, with 1.5X optical zoom on a Nikon Eclipse Ti inverted microscope controlled using Elements software). Cells were maintained at 37°C with 5% CO_2_ using a Tokai Hit stage incubator. Metaphase cells were identified using the appearance of a tight metaphase plate using DIC. The needle was then positioned close to the cell until the pipette made firm contact with the cell. Negative pressure was then applied using a Fluigent microfluidic pressure control system (Flow EZ) controlled through Fluigent controller software on the microscope computer. Frames were acquired every 2 seconds, with pressure increased by 0.5 mbar every 10 seconds. Cortex tension was then calculated by analyzing the DIC time montages using a custom MATLAB script. Tension (T) was calculated as

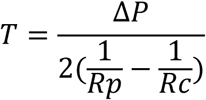

where, ΔP= pressure difference, R_p_= radius of pipette, R_c_= radius of cell.

### Calibration of protein expression using immunofluorescence

Calibration of protein expression was performed using an immunofluorescence based approach as described previously (Supplementary Fig. 7) (Taneja and Burnette, 2019). siRNA resistant MII was introduced into the knockdown cells using transient transfection, plated on glass coverslips, followed by fixation. Untransfected scrambled control cells were prepared in parallel. The two samples were then stained for endogenous MIIA or MIIB in the red channel. The samples were then imaged using the same parameters for the red and green channels in growth medium at 37°C as performed for the actual experiment. GFP intensity measured in the cytoplasm was then calibrated against the endogenous (red) signal relative to the scrambled control. The intensity in the cytoplasm was also normalized against relative enrichment in the cytoplasm as well as background. This analysis was performed for every different objective lens used in the study. Since we used the same imaging parameters for GFP throughout the experiments this allowed us to compare levels to endogenous.

### Data Quantification

For quantification of MII ensemble length (Fig. 1A-C), a 15 µm by 8 µm region of interest containing the cleavage furrow was cropped from maximum projected SIM micrographs. Only bright, in-focus, objects were included in the analysis. Measurements were made manually with individual lines saved as ROIs in Fiji and then exported for calculation of average ensemble length. Separate investigators acquired and analyzed the data for ensemble length quantification. The investigator analyzing the data was blinded.

For measuring the rate of cleavage furrow ingression, phase contrast or DIC time montages were recorded at 30 second intervals. Individual montages were cropped and aligned using the StackReg plugin in Fiji. Kymographs were created using the Multiple Kymograph plugin with a 3 pixel thick line drawn across the equator. Ingression rates were calculated as the change in distance (in µm) divided by the amount of time (in min) between the start and end of ingression (Fig. 1D).

To quantify blebbing, DIC time-lapse recordings at 30 second intervals were used. We counted blebs between the time point immediately following the end of Anaphase A (sister chromatids no longer move away from each other; this was also verified by making a kymograph across the long axis of the cytokinetic cell) and the end of cleavage furrow ingression. Every bleb that was in focus and could be accurately measured was counted for quantification. The total surface area for all blebs counted was calculated assuming blebs to be spherical. This was then normalized to the number of minutes for which the analysis was performed.

For quantification of binucleate cells, a large stitch of cells stained with phalloidin and DAPI was acquired using a 20X objective (shown in Supplementary Fig. 1C or smaller field of view shown in in Fig. 2A). Binucleate cells were manually counted using the multi-point tool in Fiji. Cells very close together in clusters and whose boundaries could not be accurately discerned were not included in the analysis.

To quantify ingression rates within hESC colonies (Supplementary Fig. 2), images were recorded at 30 second intervals. Distance within the colony was measured in “cell radii” from the edge. The colonies were assumed to be composed of concentric circular cell layers increasing in number from the edge. For any given dividing cell, its position was determined by counting the smallest number of cell radii to reach the edge. Ingression rates were measured using the MultipleKymograph plugin in Fiji. Kymographs where the cell boundary could not be ascertained were discarded.

To quantify cell shape changes upon MII knockdown, we measured the increase in pole-to-pole elongation during cytokinesis relative to metaphase. A line was drawn across the long axis of the cell from phase contrast time montages. We observe that the distance between the cell boundary during metaphase is smallest immediately before the onset of anaphase, which is in agreement with the general idea that maximal de-adhesion must occur for cell-cycle progression (Jones et al., 2018; Marchesi et al., 2014). Following anaphase onset, pole-to-pole expansion was observed, until ingression is completed, which was then followed by poles moving closer together again prior to cell spreading. We measured the pole-to-pole distance immediately upon anaphase onset (as shown in schematic, Fig. 4B), and the maximal pole-to-pole distance before they come closer together (as shown in schematic, Fig. 4B). This was also visually confirmed in the live cell montages. These widths were used to calculate the pole-to-pole elongation ratio.

To generate the endogenous polar cortex and equatorial cortex localization profiles upon MII paralog knockdown (cf. Fig. 6A-C), cells were grouped into early (>20 µm), mid (6-20 µm) and late cytokinesis (<6 µm) based on degree of furrow closure, as done previously (Taneja et al., 2016). Polar cortex intensity was calculated as the mean fluorescence averaged from 6 ROIs (1.8 µm^2^ area) placed in the same positions (see inset schematics, Fig. 6). Due to the non-uniform distribution of cortical signals, this approach allowed us to get an unbiased measurement of mean cortical intensity at the polar cortex. Equatorial cortex intensity was calculated as mean fluorescence of 2 ROIs (1.8 µm^2^ area) placed on each side of the furrow. Mean equatorial or polar cortex intensities were then averaged within the cytokinesis groups. Any perturbations were normalized to the mean of the control data sets.

### Statistics

Statistical significance for averaged data from multiple experiments, which is always depicted using bar graphs, was determined using unpaired, 2-tailed, homoscedastic Student’s T-test performed on Excel. For data pooled from multiple experiments, always depicted as Tukey plots, Mann Whitney U-test was performed on GraphPad Prism. Error bars in all bar graphs represent standard error of the mean, while Tukey plots were represented with boxes (with median, Q1, Q3 percentiles), whiskers (minimum and maximum values within 1.5 times interquartile range) and outliers (solid circles). No outliers were removed from the analysis.

For graphs displaying percentages, no error bar was displayed and the data over more than 3 independent experiments was pooled. Due to the pooled approach for the endogenous MII localizations in HeLa cells, error bars depict standard error of weighted means, where the weights were determined based on how many cells from a particular experimental replicate were allocated to a particular group (e.g. Early, Mid, Late). The weighted standard deviation was calculated to remove any bias from individual experiments, using the formula 

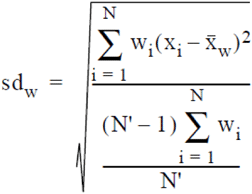

where w_i_ is the weight for the i^th^ observation, N’ is the number of non-zero weights, and x_w_ is the weighted mean of the observations

## SUPPLEMENTARY INFORMATION

**Supplementary figure 1.**
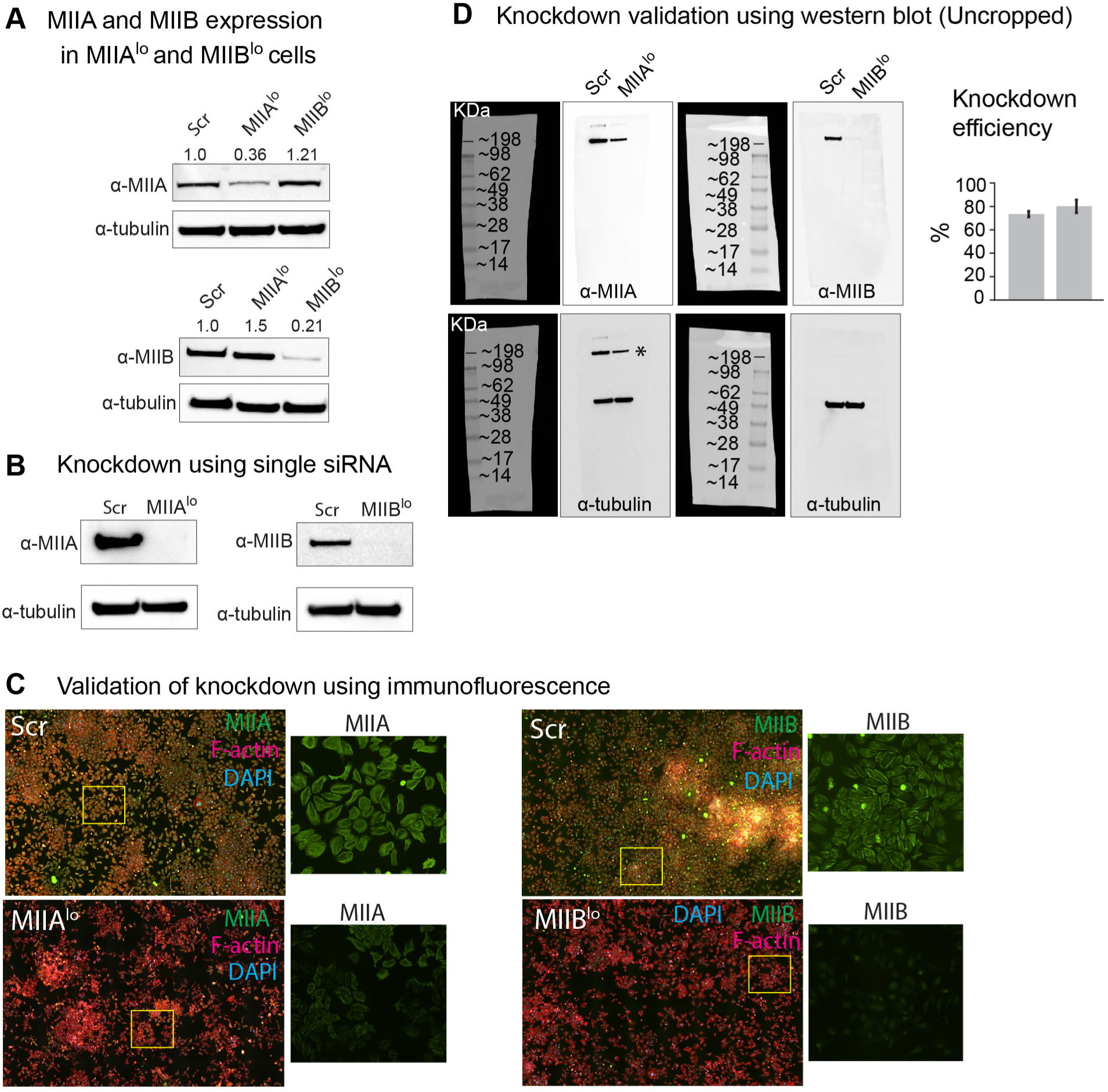
Characterization of MII paralogs in HeLa cells. A) MIIA and MIIB expression upon depletion of the other paralog. Tubulin was used as loading control. The numbers represent expression normalized to loading control with scrambled control set to 1. B) Knockdown of MIIA and MIIB using individual siRNA. The numbers represent Mil band intensity normalized to loading control. C) Large stitches of Scr, MIIA^lo^ and MIIB^lo^ cells stained for F-actin, DAPI and the depleted isoform. Insets: Enlarged view of yellow box. D) Uncropped representative western blot from another experiment validating MIIA and MIIB knockdown. Tubulin was used as loading control for normalization and calculation of knockdown efficiency. N=5 independent knockdowns. *, leftover MIIA bands from blot stripping. Error bars represent standard error of the mean.

**Supplementary Figure 2.**
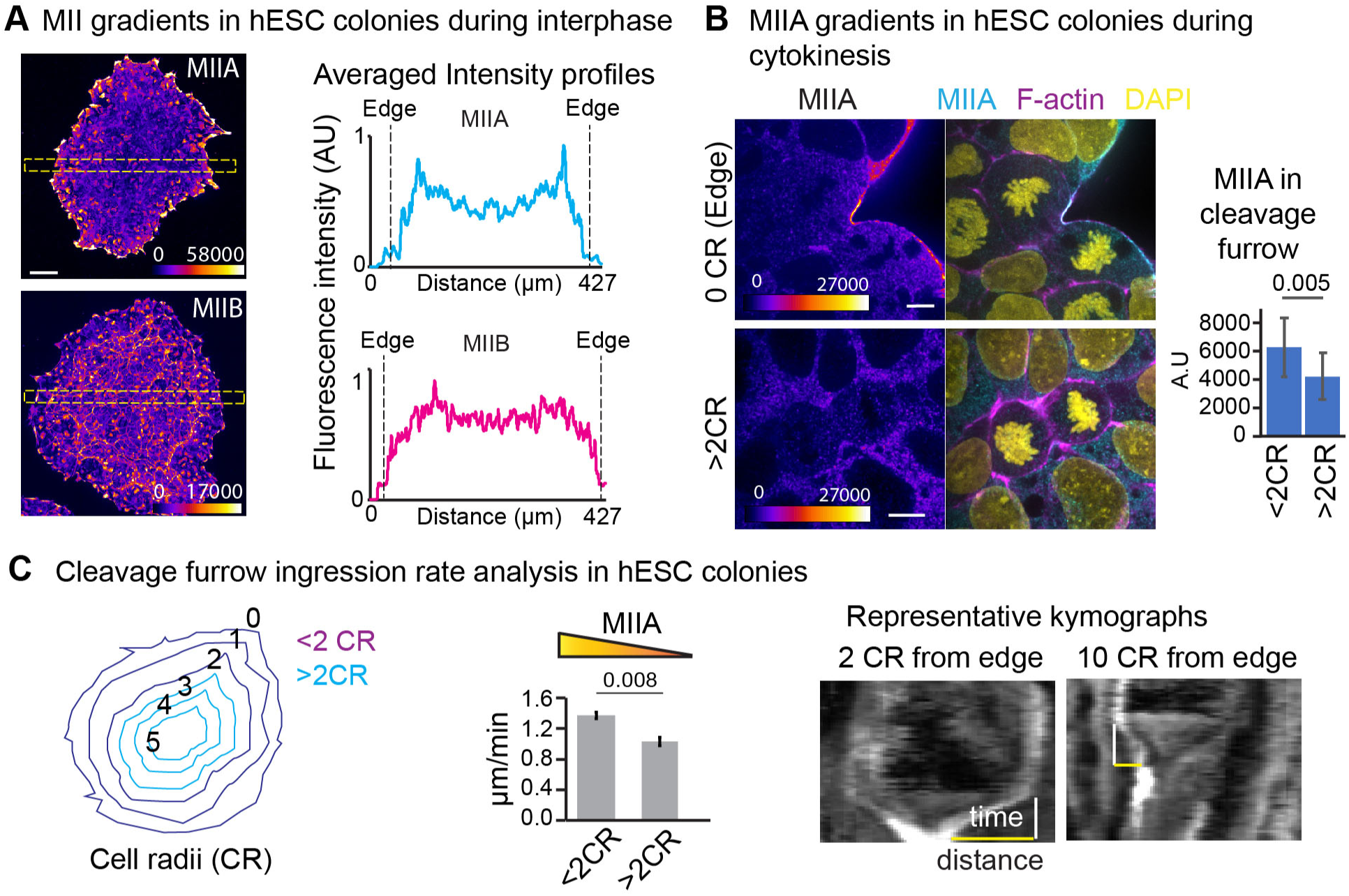
MIIA drives faster ingression in human embryonic stem cell colonies. We sought a system where natural gradients of MIIA exist, for which we turned to hESC colonies, that show a gradient of MIIA with high MIIA at the edge and less in the centre. MIIB was showed no gradient in localization. We used this system to test whether MIIA levels correlate with faster ingression. A) hESC colonies stained for endogenous MIIA or MIIB (Fire LUT). Line scans from yellow box averaged over 12 30° rotations. B) Representative dividing cells and expression of MIIA in the cleavage furrow of dividing cells close to (<2CR) or away from (>2CR) the edge of a hESC colony. Intensity matched micrographs show endogenous MIIA staining represented using Fire LUT. Quantification from 14 <2CR and 17 >2CR cells from 10 separate colonies. C) Representative kymographs from phase contrast time montages of hESCs dividing 2 CR or 10 CR from the edge of the colony and cleavage furrow ingression rates of cells dividing in hESC colonies. n=61 cells from 5 colonies over 3 independent experiments. Cell position was represented as cell radii (CR) away from the edge assuming a spherical colony with each concentric layer of cells spanning 1 cell radius (see inset schematic). Scale bar in (A) and (D)- 50 µm and 10 µm, respectively. Error bars in (C) represent standard error of the mean; Error bars in (B) represent standard deviation. p values depicted over respective bar graphs.

**Supplementary Figure 3.**
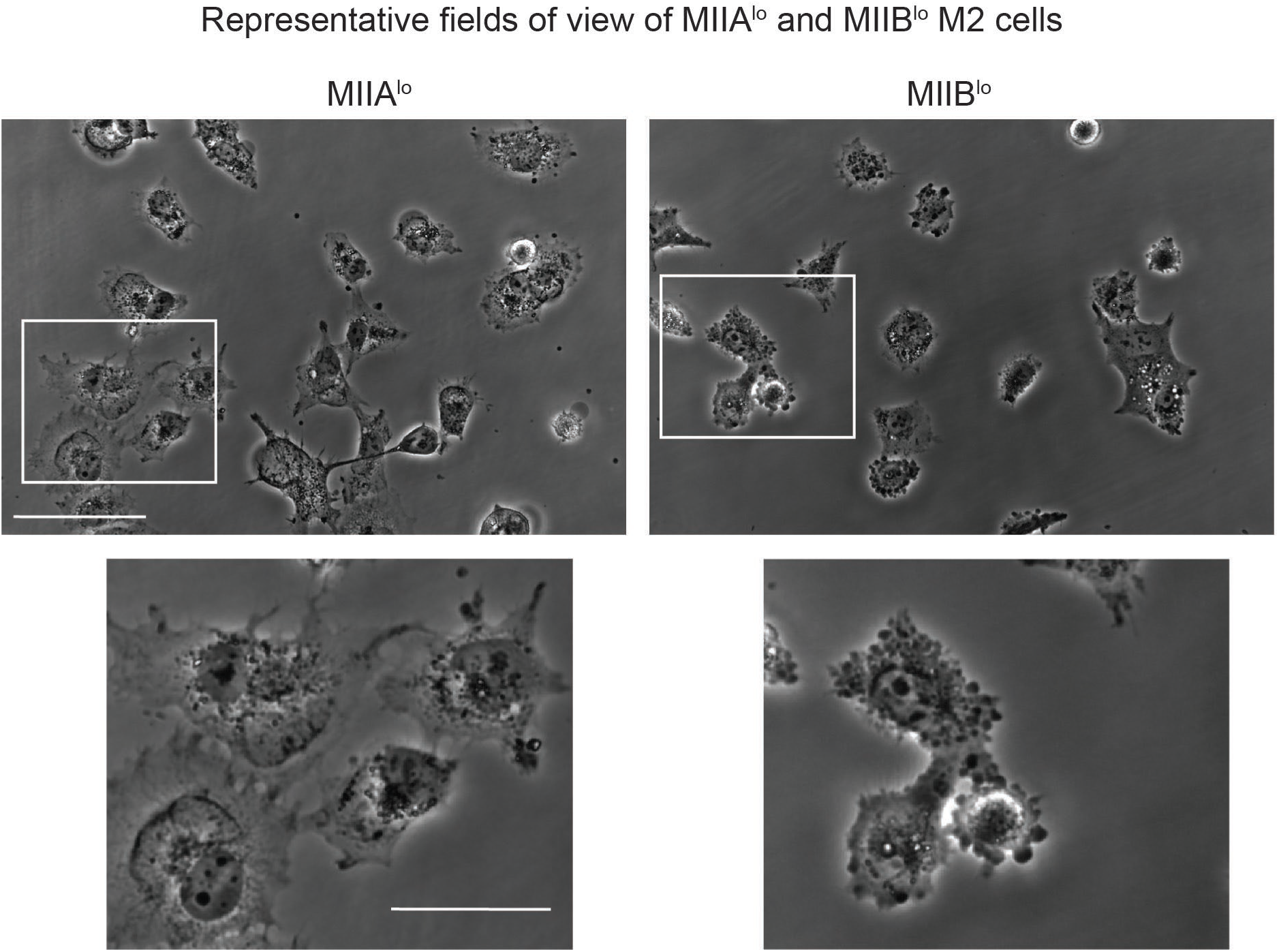
Depletion of MIi paralogs in M2 cells. Representative fields of view of MIIA^lo^ and Mll8^lo^ M2 cells at 5 hours post plating. Insets-Enlarged views of white box. Scale bar- 100 µm

**Supplementary Figure 4.**
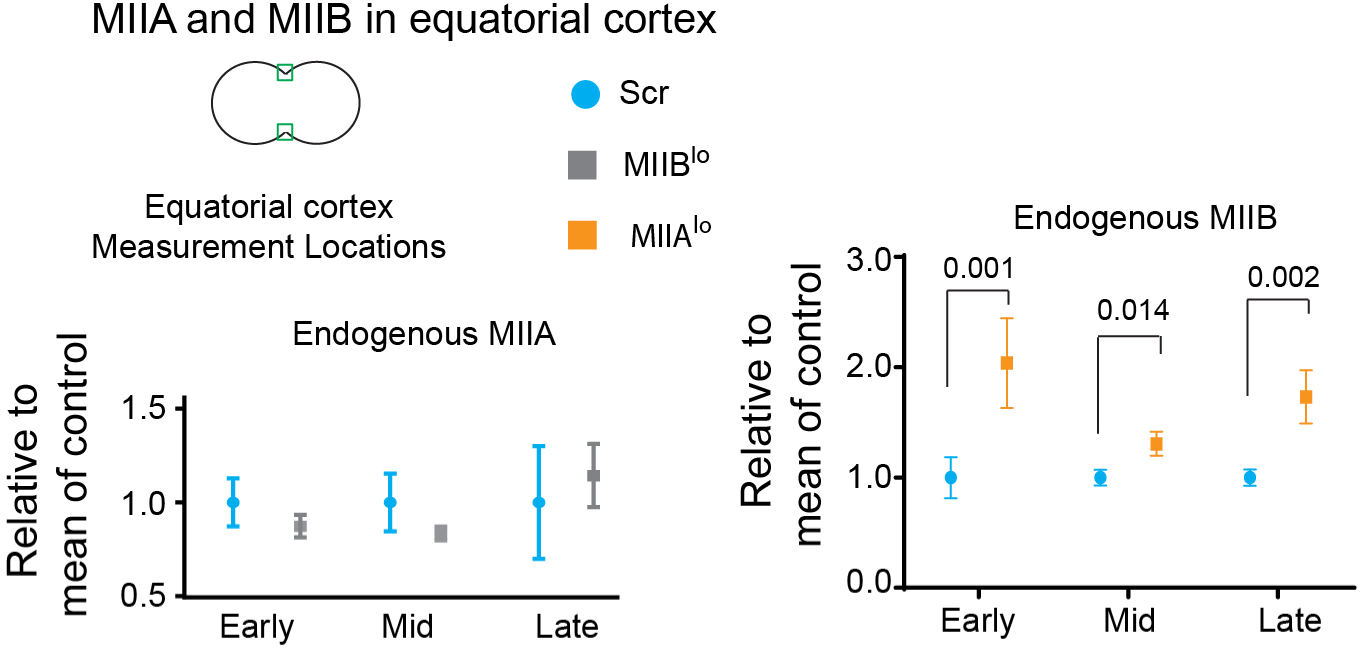
MIi paralog compensatory localization during C-phase. Quantification of MIIA and MIIB at the equatorial cortex. Intensity was calculated as mean of green ROls in cartoon insets. See methods for detailed quantification scheme. For MIIA: n=41 Ser cells and 48 MIIB^10^ cells over 3 independent experiments. For MIIB: n= 43 MIIA^lo^ and 46 Ser cells over 3 independent experiments. Error bars represent standard error of mean.

**Supplementary Figure 5.**
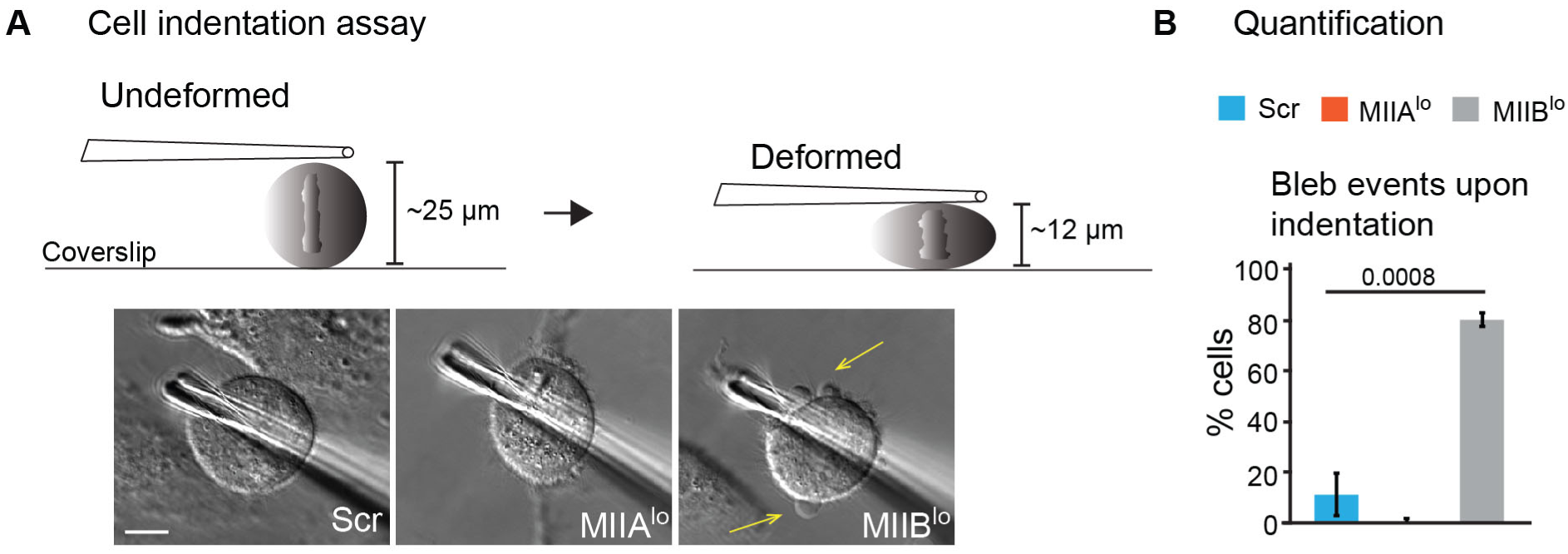
Cell indentation assay. A) Cell indentation assay design. DIC montage of Ser, MIIAlo and MIiBio cells upon indentation. B) Quantification of blebbing events. n=16 Ser cells, 11 MIIA^lo^ and 21 MIIB^lo^ cells over 3 independent experiments. Scale bar- 10 µm. Error bars represent standard error of the mean. p values stated over respective bars.

**Supplementary Figure 6.**
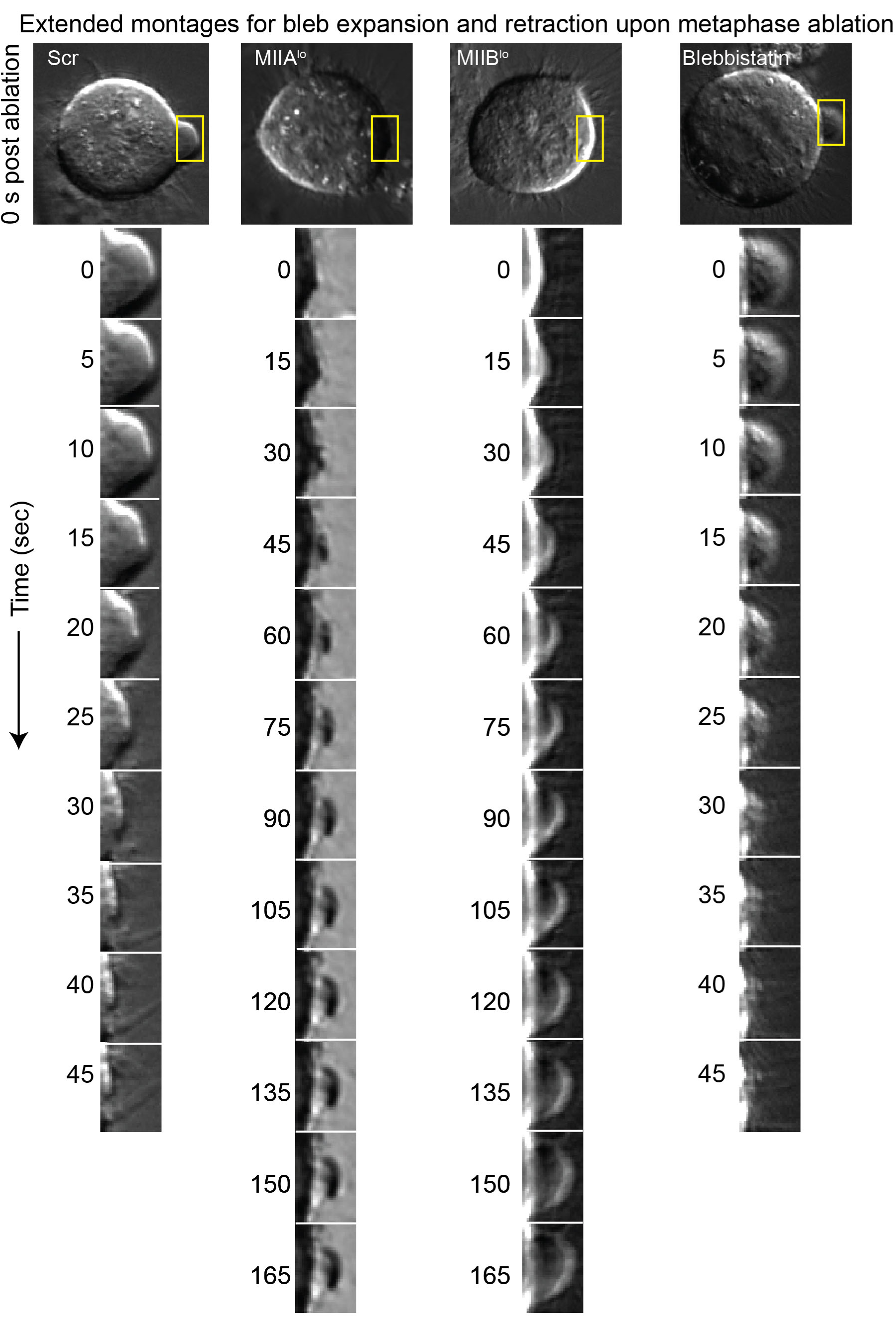
Extended montages for metaphase ablation upon MII depletion. Montage corresponds to yellow box.

**Supplementary Figure 7.**
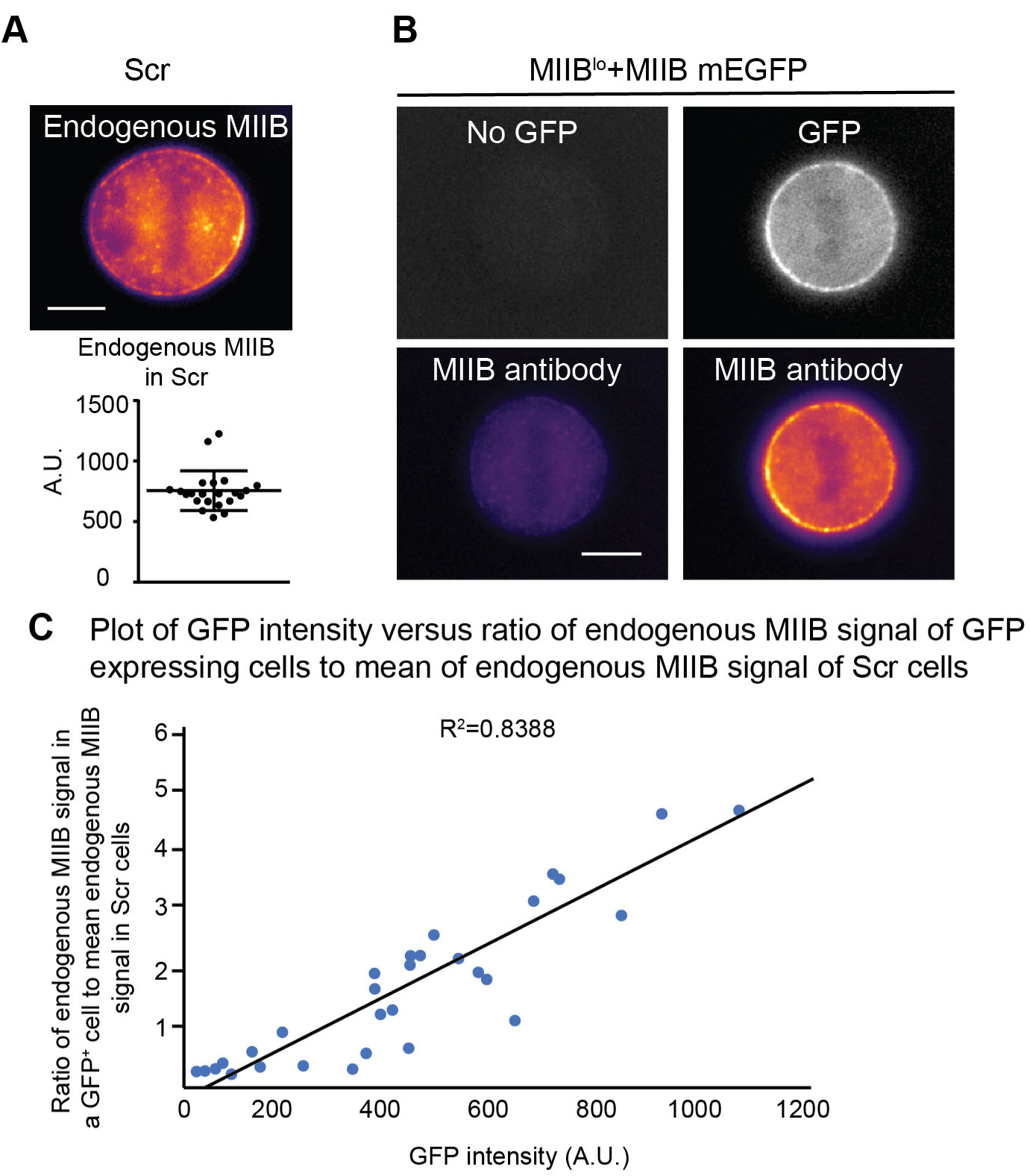
Calibration of protein expression using immunofluorescence. Representative example of endogenous MIIB localization in Ser cells. Graph shows the distribution of cortical signals measu red using 21 cells. Error bars show standard deviation. See methods for detailed quantification method. B) Two examples of endogenous MIIB localization in MIIB^10^ cells re-expressing full length MIIB-mEGFP. In the first example (left), a cell with no GFP expression is shown. In the second example, a cell expressing GFP is shown. Endogenous MIIB is colored using mpl-inferno LUT. _C)_ Using several cells expressing varying levels of GFP, the GFP signal was plotted against the ratio of the endogenous MIIB signal of GFP expressing cells to the mean of the endogenous MIIB intensity in Ser cells (plotted in (A)). By using the same imaging parameters for every experiment, this allowed us to extrapolate the expression in our rescue experiments with respect to wild type levels using the linear fit of the plot above. Scale bar- 10 µm.

**Table S1.**
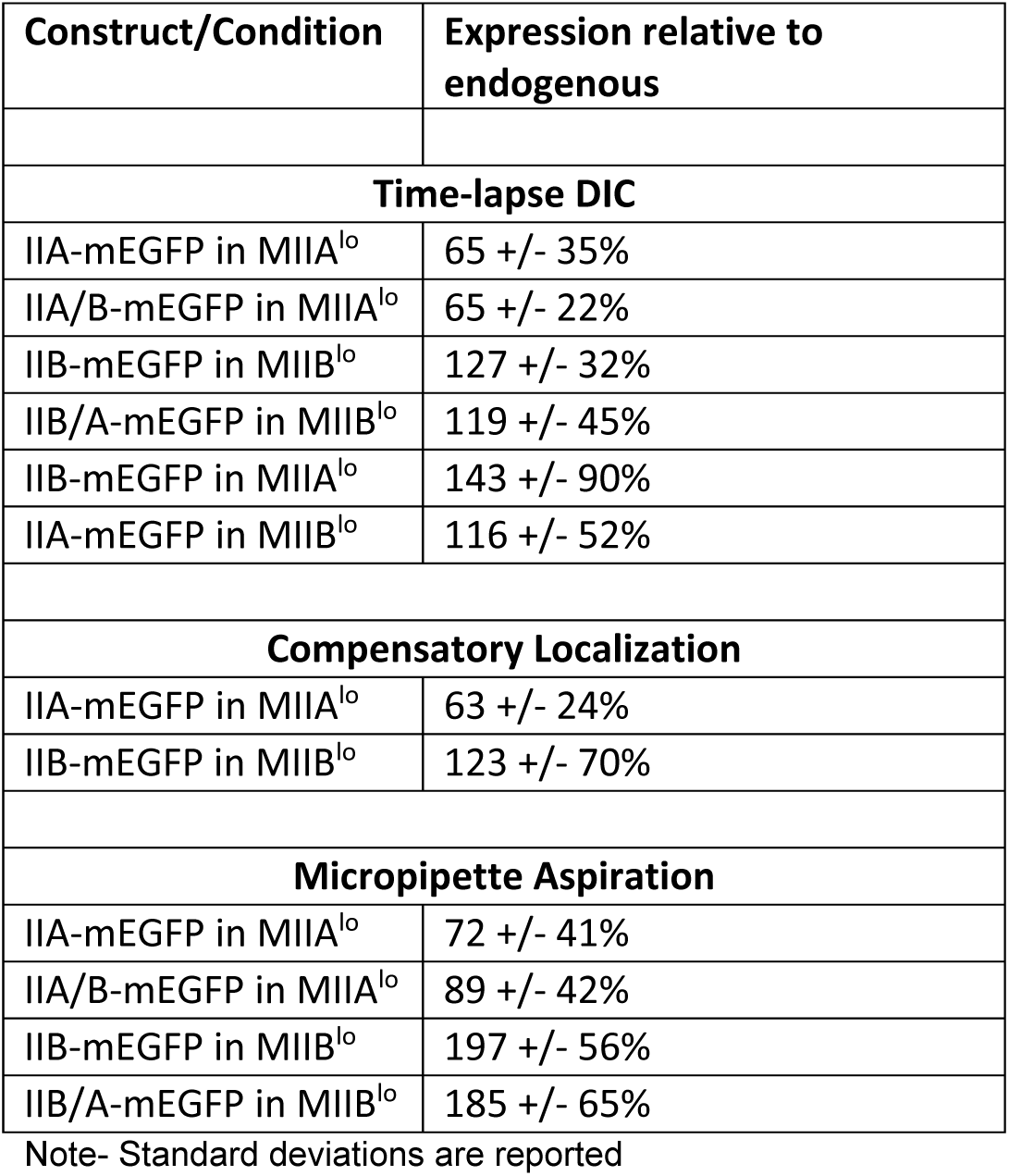

## Movie Legends

**Supplementary Movie 1**. Time lapse DIC imaging of control HeLa cell undergoing cytokinesis. Note modest blebbing during furrow ingression. Movie dimensions: 82 µm by 52 µm. Length: 24.6 min.

**Supplementary Movie 2**: Time lapse DIC imaging of MIIB depleted cell undergoing cytokinesis. Note dramatic blebbing and spindle rocking during furrow ingression. Movie dimensions: 82 µm by 52 µm. Length: 31.8 min.

**Supplementary Movie 3**: Time lapse DIC imaging of MIIA depleted cell undergoing cytokinesis. Note complete absence of blebbing during furrow ingression. Movie dimensions: 82 µm by 52 µm. Length: 29.2 min.

**Supplementary Movie 4**: Time lapse confocal imaging of MIIA depleted HeLa cell expressing MIIB mEmerald undergoing cytokinesis. Movie dimensions: 81 µm by 31 µm. Length: 34 min. Movie dimensions: 49 µm by 45 µm. Length: 33.1 min.

**Supplementary Movie 5**: Time lapse confocal imaging of MIIB depleted HeLa cell expressing MIIA mEmerald undergoing cytokinesis. Note the onset of blebbing is correlated with the enrichment of MIIA at the cortex. Movie dimensions: 81 µm by 31 µm. Length: 34 min. Movie dimensions: 49 µm by 45 µm. Length: 29 min.

**Supplementary Movie 6**: Time lapse confocal imaging of HeLa cell expressing MIIA mApple (red) and MIIB mEmerald (green) undergoing cytokinesis. Note enrichment of MIIB at polar cortex prior to ingression. MIIA and MIIB are both enriched first at the equatorial cortex followed by the polar cortex. Movie dimensions: 81 µm by 31 µm. Length: 34 min.

**Supplementary Movie 7**: Time lapse DIC imaging of control cell indented using micropipette to half its height. Movie dimensions: 65 µm by 61 µm. Length: 6.9 min.

**Supplementary Movie 8**: Time lapse DIC imaging of MIIB depleted cell indented using micropipette to half its height. Note blebbing induced upon indentation. Movie dimensions: 65 µm by 61 µm. Length: 6.9 min.

**Supplementary Movie 9**: Time lapse DIC imaging of MIIA depleted cell indented using micropipette to half its height. Movie dimensions: 65 µm by 61 µm. Length: 6.9 min.

